# Doublecortin engages the microtubule lattice through a cooperative binding mode involving its C-terminal domain

**DOI:** 10.1101/2021.02.17.431714

**Authors:** Atefeh Rafiei, Linda Lee, D. Alex Crowder, Daniel J. Saltzberg, Andrej Sali, Gary J. Brouhard, David C. Schriemer

## Abstract

Doublecortin (DCX) is a microtubule (MT) associated protein that regulates MT structure and function during neuronal development and mutations in DCX lead to a spectrum of neurological disorders. The structural properties of MT-bound DCX remain poorly resolved. Here, we describe the molecular architecture of the DCX-MT complex through an integrative modeling approach that combines data from X-ray crystallography, cryo-EM and a high-fidelity chemical crosslinking method. We demonstrate that DCX interacts with MTs through its N-terminal domain and induces a lattice-dependent self-association involving both the C-terminal structured domain and the C-tails, in a conformation that favors an open, domain-swapped state. The networked state can accommodate multiple different attachment points on the MT lattice, all of which orient the C-tails away from the lattice. As numerous disease mutations cluster in the C-terminus, and regulatory phosphorylations cluster in the C-tail, our study shows that lattice-driven self-assembly is an important property of DCX.

## Introduction

The regulation of microtubule (MT) polymerization is central to the maintenance of cell polarity, intracellular transport and cell division. Aside from the intrinsic dynamics of the MT itself, polymer nucleation, growth and disassembly are coordinated by many different microtubule-associated proteins (MAPs) to establish and manage the wider cytoskeleton in a cell-type dependent manner. One significant promoter of MT regulation in developing neurons is the X-linked *doublecortin* gene. Doublecortin (DCX) is a 40 kDa MAP, essential for neuronal migration in embryonic and postnatal brain development (des Portes et al., 1998a, Gleeson et al., 1998, Francis et al., 1999). DCX influences MT rigidity and curvature and selects for the canonical 13-protofilament geometry that is characteristic of *in-vivo* MTs (Moores et al., 2004, Jean et al., 2012, Bechstedt and Brouhard, 2012). It also increases the nucleation rate and decreases the catastrophe rate (Moores et al., 2006), producing a net stabilization of MTs.

Mutations in the DCX-encoding gene result in lissencephaly (“smooth brain”) or subcortical band heterotopia, both of which can generate a spectrum of intellectual inabilities and/or seizures (des Portes et al., 1998b). These mutations impair binding of DCX to microtubules (Taylor et al., 2000) but the structural basis for this effect remains unclear. DCX consists of a tandem repeat of Doublecortin-like (DC) domains possessing 27% sequence identity, an N-terminal tail (N-tail), a linker region, and a long Serine/Proline-rich C-terminal tail (C-tail) (Burger et al., 2016a). Both of the DC domains and the linker region are necessary for the MT stabilization effect (Taylor et al., 2000, Kim et al., 2003), whereas the CDC (Manka and Moores, 2020) and the C-tail (Moores et al., 2004) help establish the preference for 13-protofilament MTs. The C-tail is also implicated in regulating microtubule-actin interactions (Fu et al., 2013, Jean et al., 2012, Tsukada et al., 2005). Structures are only available for the isolated N-terminal DC domain (NDC) (Kim et al., 2003, Cierpicki et al., 2006, Burger et al., 2016a) and the C-terminal DC domain (CDC) (Burger et al., 2016a, Rufer et al., 2018). For the most part, these structures share the same globular ubiquitin-like fold, although the CDC may adopt a substantially opened form (Rufer et al., 2018). Disease mutations cluster in both of the structured domains, suggesting a functional conservation based upon a shared property, possibly a direct interaction with the MT lattice (Taylor et al., 2000).

Cryo-Electron Microscopy (cryo-EM) investigations of the DCX-MT interaction reveal that one of the two structured domain binds to MTs at the junction of four α/β-tubulin dimers, stabilizing the longitudinal protofilament and its lateral contacts (Fourniol et al., 2010, Liu et al., 2012). Given the modest resolution of the cryo-EM maps and the structural similarity of DC domains, there is still debate over the identification of the actual interacting DC domain. NDC was designated as the primary contact at the junction, but evidence also suggests that CDC could occupy this site (Burger et al., 2016a). The binding mode is clouded further by observations that DCX has a propensity to dimerize *in vitro* (Caspi et al., 2000), although analytical ultracentrifugation showed that isolated DCX is monomeric even at relatively high concentrations (Moores et al., 2006). However, the C-terminal domain may adopt an open conformation, allowing it to dimerize through a “domain swapping” event in some MT-assisted fashion (Rufer et al., 2018). A recent study describes a dynamic interaction model in which DCX interacts with the MT through the CDC domain first, then transitions to an NDC-MT binding mode in the fully assembled MT lattice (Manka and Moores, 2020). The various models presented are incompatible. It is difficult to rationalize how a dimerizing DCX could interact with the MT lattice in a manner that allows for both NDC and CDC binding and explain the density observed in the cryo-EM measurements.

Greater clarity on the primary modes of MT engagement would help address the structural basis for the observed mutational effects, as well as help resolve additional structure-function problems. For example, DCX manifests a higher affinity to MT ends compared to the MT lattice (Bechstedt et al., 2014) and appears sensitive to MT lateral (Bechstedt and Brouhard, 2012) and longitudinal curvature (Bechstedt et al., 2014). These observations are difficult to explain through a simple binding mode, particularly given that binding is strongly cooperative and significantly reduces the rate of MT dissociation (Bechstedt and Brouhard, 2012). Further, how DCX is functionally altered by phosphorylation (Tanaka et al., 2004, Graham et al., 2004, Shmueli et al., 2006) and engages coregulatory proteins like Neurabin II (Tsukada et al., 2005) and kinesin-3 motor protein (Liu et al., 2012, Lipka et al., 2016) requires a complete structure of the full length protein on the MT lattice. Here, we develop a unifying model of the DCX-MT interaction through an integrative structure-building approach, using data from an improved crosslinking mass spectrometry (XL-MS) method (Ziemianowicz et al., 2018, Rafiei and Schriemer, 2019), together with available cryo-EM and X-ray crystallographic structures. Our results support a DCX-MT interaction model in which NDC binds to MTs at the junction of the four α/β-tubulin dimers, and induces DCX self-association through its CDC and C-tail domains, creating an extended structure capable of capturing different lateral and longitudinal MT interactions.

## Results

### Preparation of the DCX-MT construct

The activity of the recombinantly purified DCX (**Fig. S1A**) was confirmed using three different methods. In a turbidity assay, adding DCX to purified α/β tubulin induced MT polymerization (**Fig. S1B**), consistent with previous reports (Bechstedt and Brouhard, 2012). The effect saturates at 10-20 µM DCX, in line with previous claims of a 1:1 binding ratio (DCX:α/β-tubulin) (Moores et al., 2006). A pelleting assay confirmed a 1:1 stoichiometry (**Fig. S1C**). Finally, using a fluorescence image analysis, we observed MT lengths to decrease with increasing DCX concentration, confirming a role in MT nucleation (**Fig. S1D**) (Moores et al., 2004). There were no signs of MT bundling (**Fig. S1E**).

### Sampling the equilibrated interaction with photoactivated crosslinking

Many different crosslinking reagents are available for measuring site-to-site distances, but most are not appropriate for structural characterization of dynamic systems. The inherent flexibility of DCX (**Fig. 1A,B**) renders it susceptible to “kinetic trapping” on the MT lattice when using conventional long-lived reagents (Ziemianowicz et al., 2018), thus potentially scrambling the sites of interaction. That is, conformations not representative of the structural ensemble can be selected based simply on higher reaction rates. Therefore, we used a crosslinker method optimized for MT interactions involving the heterobifunctional reagent LC-SDA that has been demonstrated to minimize this effect (Ziemianowicz et al., 2018, Rafiei and Schriemer, 2019). The first coupling is to accessible nucleophiles through a conventional NHS ester and the second coupling to any surface-accessible residue through laser-initiated carbene chemistry. When applied to the saturated DCX-MT state, the method generated a dense set of 362 unique crosslinks, well-distributed among the domains and subunits (**Fig. 2A** and **Table S1**), including 124 inter-protein crosslinks between DCX and α/β-tubulin.

**Fig. 1.**
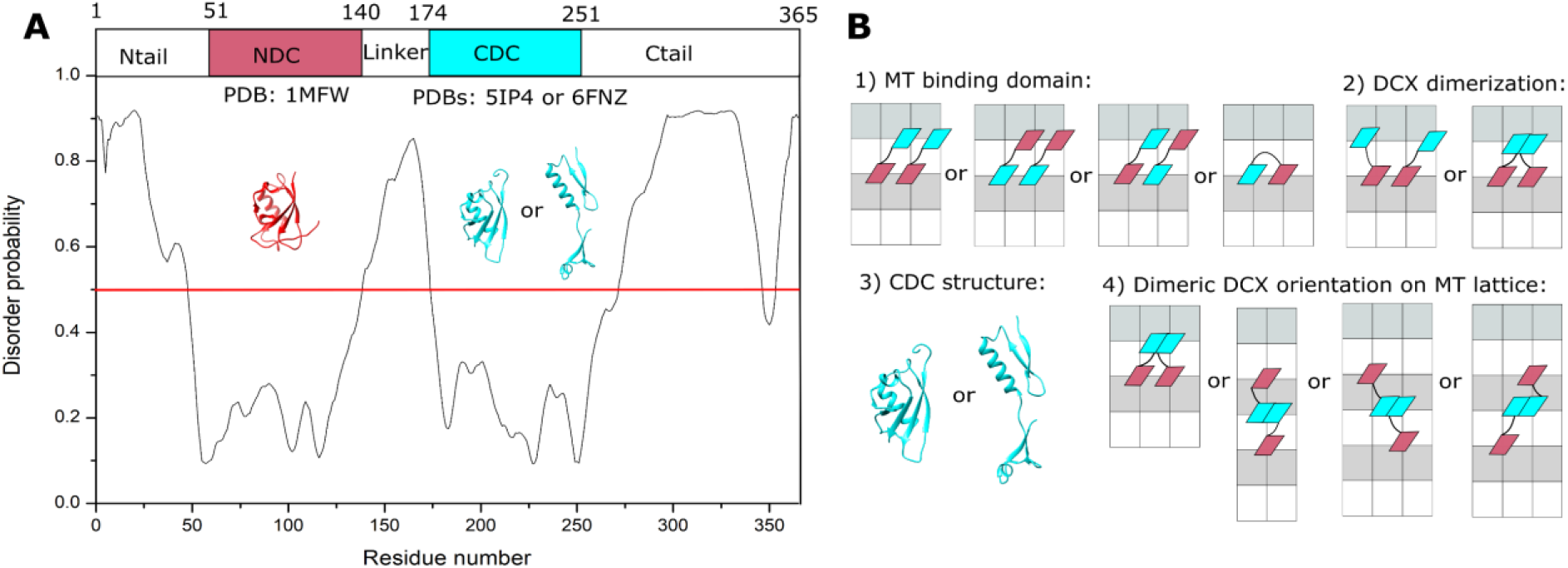
DCX structure and lattice interaction options. **A)** The disordered regions of DCX sequence predicted using PrDOS (Ishida and Kinoshita, 2007). B) Schematic representations of the four orientational challenges to the elucidation of the DCX-MT binding mode. In each case, a minimal DCX construct is shown, comprising NDC (red), linker (black) and CDC (cyan). α−Tubulin and β−tubulin are shown with white and light grey rectangles, respectively.

**Fig. 2.**
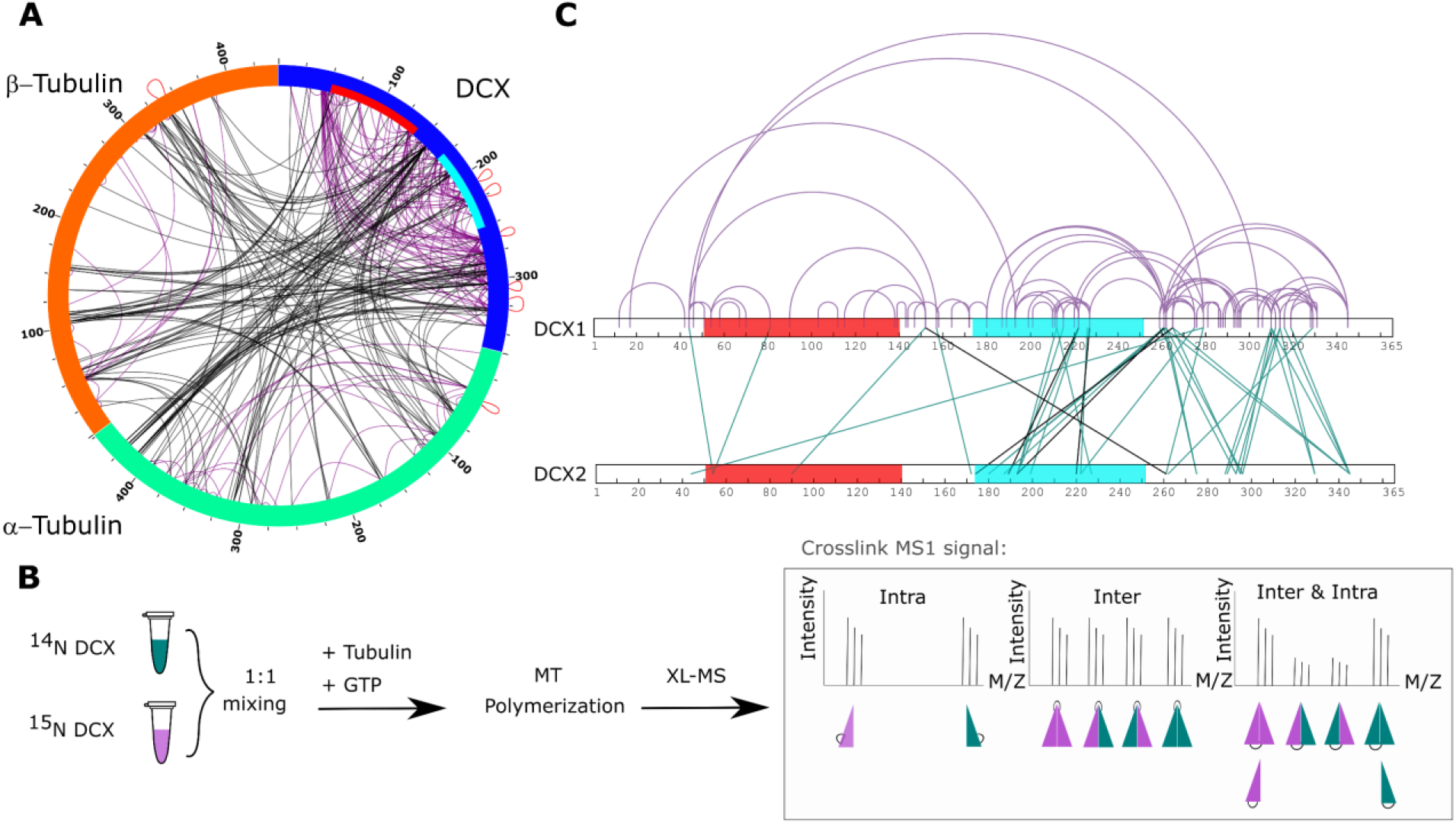
Crosslinking-mass spectrometry analysis of DCX. **A)** Two-dimensional crosslinking map linking α−β Tubulin and DCX at specific residues. NDC and CDC domains of DCX sequence are shown with red and cyan, respectively. Intra-protein crosslinking sites are shown in black and and inter-protein crosslinks in purple. A subset of inter-protein crosslinking sites observed between peptides with a shared sequence are shown in red loops. The crosslinking map is produced using xVis (Grimm et al., 2015). **B)** Inter- and intra-DCX crosslinking sites differentiated using a protein isotopic labeling technique. Three categories are defined, where the ratios of intensities for the inner and outer doublet in the mixed state are variable. The crosslinking map is produced using xiNET online tool (Combe et al., 2015). **C)** Two-dimensional crosslinking map linking DCXs at specific residues. NDC and CDC domains of DCX are shown in red and cyan, respectively. Shared inter and intra XLs are shown as inter (green) and intra (purple) DCX crosslinks. Unique inter-DCX crosslinking sites are shown in black. In all cases, when there is ambiguity in the crosslinking site, a single crosslinking site pair was used for visualization.

These datasets were then used for integrative structure determination. Given the multiplicity of binding modes that are possible between DCX and the MT lattice, we reasoned that an incremental approach to modeling focused on determining the major interaction modes would be necessary for computational efficiency. Thus, we formulated the modeling exercise into four stages (**Fig. 1B**): (1) identify the primary binding domain using only crosslink restraints between MT and structured DCX domains, (2) evaluate if MT engagement induces DCX dimerization using inter-DCX crosslink restraints, (3) determine if CDC can adopt a domain-swappable (open state) conformation using only DCX crosslinks, and finally (4) determine the orientation(s) of DCX on the MT lattice, constrained by major modes determined in 1-3 and the full set of crosslinking data. These modeling stages were informed with available crystal structures, cryo-EM maps, and the crosslinking mass spectrometry restraints as required, using a four-step workflow (Rout and Sali, 2019, Alber et al., 2007, Webb et al., 2018) (**Fig. S2**).

### NDC is the main MT binding domain

A minimal MT “repeat unit” was established for modeling, consisting of 2 α−tubulin and 4 β−tubulin subunits. This repeat unit contains all possible lateral and longitudinal tubulin-tubulin interactions found in the main lattice-B state. As DCX is excluded from lattice-A interactions (Fourniol et al., 2010, Manka and Moores, 2018), this lattice type was not built into our model representation. The NDC and CDC domains were then tested separately for their occupancy of the major binding site. Integrative modeling was performed in two ways: first using only crosslinks, and then crosslinks combined with the available DCX-MT EM density map (Liu et al., 2012). This approach allowed us to determine, by comparison, how well the crosslinking data alone could locate the expected binding site and thus validate the quality of the crosslinks.

Two major clusters of solutions were obtained for each domain when using the crosslinks alone (**Fig. S3** and **Table S2**). For NDC-MT, the top cluster contained 48% of all models (cluster precision of 23.4 Å) and identified a site at the junction of the four tubulin dimers resembling the site identified by cryo-EM, albeit with a slightly altered orientation (**Fig. 3A,E**). The most accurate models, as measured by RMSD from the expected site also had the highest crosslink utilization rate (**Fig. 3A**). The next major cluster, containing 36% of models (cluster precision of 25.0 Å), was identified at the partial binding site at the edge of the repeat unit (**Fig. S4A**). The top cluster for CDC-MT contained 50% of the individual models (cluster precision 27.0 Å) and identified a different location, on the inter-protofilament junction of two α−tubulin subunits (**Fig. 3B** and **Table S2**). These models are diffusive in their accuracy and crosslink utilization rate (**Fig. 3B**). The second cluster, containing 23% of models (cluster precision of 30.0 Å), was identified in the lumenal region of the MT lattice (**Fig. S4B**).

**Fig. 3.**
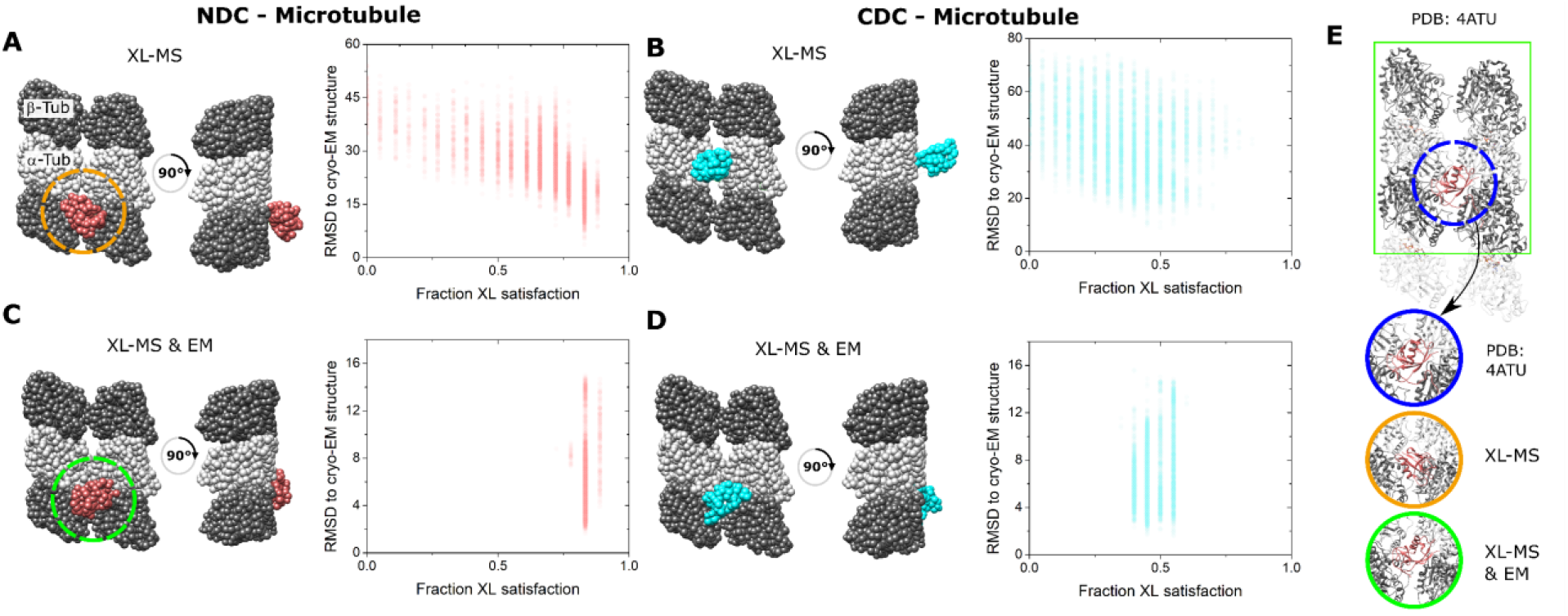
Integrative structural modeling for NDC/CDC -MT. **A)** The centroid structure for the main cluster of models produced by IMP for NDC-MT guided exclusively by XL-MS restraints. B) The centroid structure for the main cluster of models produced by IMP for CDC-MT guided exclusively by XL-MS restraints. C) The centroid structure for the main cluster of models produced by IMP for XL-MS restraints combined with the EM density map. D) The centroid structure for the main cluster of models produced by IMP CDC-MT guided by XL-MS restraints and combined with the EM density map. The fractional crosslink satisfaction (defined as <35 Å) versus RMSD to the canonical binding site (PDB 4ATU) for all models in the main structural cluster is presented for each modeling scenario. α−Tubulin and β−Tubulin are shown as light and dark grey, respectively. NDC is shown in red, and CDC in cyan. **E**) Expansions of the NDC orientations from modeling, compared to cryo-EM based NDC-MT structure (PDB 4ATU).

The addition of the EM data to the integrative modeling input, not surprisingly, returned the same binding site for NDC-MT (**Fig. 3C**). A single distinct cluster was generated that contained 80% of all models (with a cluster precision 6.0 Å) and it corrected the orientation of the domain (**Fig. 3E**). Conversely, the addition of the EM data forced the relocation of CDC to the major binding site, well removed from the one generated by crosslinking alone. Although 58% of all models clustered with a precision 5.9 Å, the crosslink utilization rate dropped considerably (**Fig. 3D**). Taken together, the high congruency between crosslinking and cryo-EM data confirms the location of NDC at the junction, whereas the variable localization of the CDC and a weaker restraint set suggests, at best, a secondary binding site. Thus, for successive stages of modeling, the NDC was located at the primary binding site and the CDC was left unrestricted.

### Doublecortin organizes on the MT lattice through CDC and C-tail domains

We next evaluated if DCX could form higher-order assemblies, and if so, which subunits are involved. To differentiate inter-subunit crosslinks from intra-subunit crosslinks, we used heavy isotopes (^15^N) installed metabolically during DCX expression. The incorporation of ^15^N was assessed by LC-MS/MS, demonstrating >99% incorporation (**Fig. S5**). Then, a 1:1 mixture of light and heavy labeled DCX (^14^N:^15^N) was used in place of light DCX in sample preparation, followed by crosslinking. Intra- and inter-protein crosslinks were differentiated based on the characteristic MS1 pattern of crosslinked peptides. We identified three types of crosslinking signatures reflective of intra-protein crosslinks, inter-protein crosslinks or a mixed state where both types can exist simultaneously although at different levels (**Fig. 2B**). This latter category is identifiable through a variable intensity pattern of the “inner doublet” (Melchior et al., 2016). We then explored if the interprotein labeling patterns could be generated without the addition of tubulin, which would indicate some measure of self-interaction in the free form, possibly dimerization or higher-order associations. We found that only under extreme cases (i.e. DCX denaturation, refolding and concentrating) could we induce a small amount of interprotein crosslinking, and only at two sites (**Fig. S6**). Thus, DCX self-association is a MT-dependent phenomenon. In total, we obtained 32 unique intra-DCX crosslink sites, 5 unique inter-DCX crosslink sites as well as 45 unique crosslink sites in the mixed group, both inter-DCX and intra-DCX (**Fig. 2C** and **Table S3**). Interestingly, >80% of the inter-DCX crosslinks identified involve the CDC domain and the C-tail, which indicates that DCX self-associates on the MT lattice through its C-terminal regions. While the inter-DCX crosslinks cannot distinguish between a dimeric state or higher-order assemblies, we chose to proceed with modeling the dimeric state as probable form of self-association based on the crystallographic model (Rufer et al., 2018).

### On-lattice dimer shows a preference for an open state

Before attempting to model the full dimeric structure using all available crosslink data, we explored if the subset of DCX-DCX crosslinks could indicate how the DCX-MT interaction might induce a dimerization event. Specifically, we performed integrative modeling to test if the data could distinguish between an interaction dominated by the globular CDC structure observed in the free form (PDB 5IP4, (Burger et al., 2016a)) or an open state reflective of a “domain-swapped” dimerization as suggested by recent crystallographic studies (PDB 6fNZ, (Rufer et al., 2018)). Two full-length DCX molecules were modeled on an expanded MT lattice to allow for all possible orientations of dimerized DCX (i.e. 3 protofilaments of 3 α/β-tubulin dimers each, **Fig. 4A**). Pairs of NDCs were placed at the confirmed junctional binding sites in four possible orientations: laterally across the protofilaments, longitudinally along the axis of the protofilaments or in one of two diagonal geometries. These multiple scenarios require assessment as the placement of the anchoring interactions could dictate the success of the dimerization event.

**Fig. 4.**
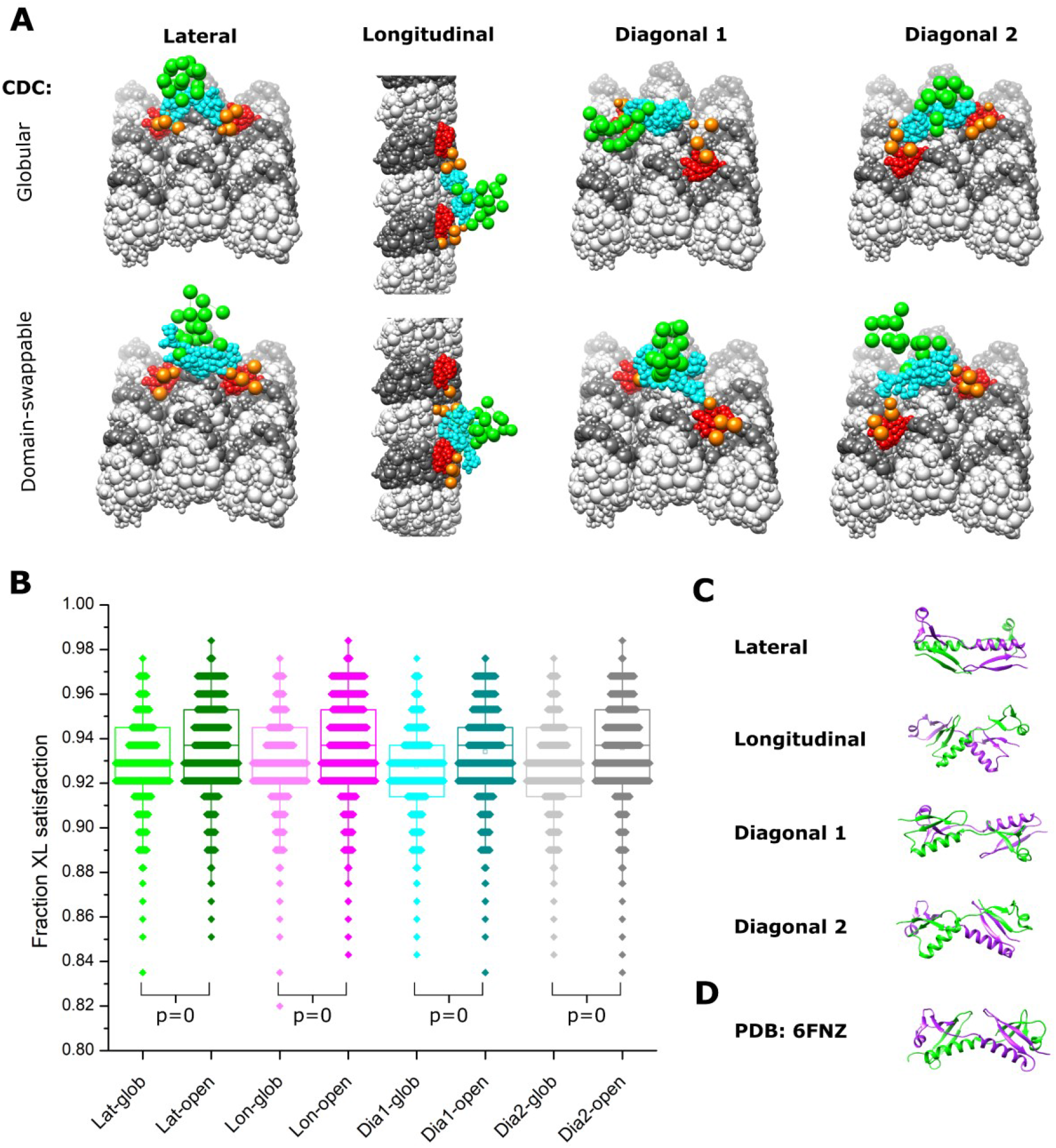
Integrative modeling of DCX self-assembly on MT lattice. **A)** The dimeric DCX-MT centroid model of the main clusters of models generated using only DCX-DCX crosslinking restraints. Four different relative positions of fixed NDC on the MT lattice were assessed (lateral, longitudinal, diagonal 1 and diagonal 2); α−Tubulin and β−Tubulin are shown as light and dark grey, respectively. NDC, linker, CDC and C-tail regions are shown as red, orange, cyan and green, respectively. CDC structure is represented as either globular or open (domain-swappable) conformations. **B)** The fractional XL satisfaction (defined as <35Å) for the main cluster of models generated for each modeling scenario. **C)** The relative orientation of the dimeric CDC structures in the centroid model of the main cluster using the domain swappable CDC conformation; green is CDC (monomer 1) and purple is CDC (monomer 2). **D**) The crystal structure of domain-swapped CDC dimer (PDB 6FNZ).

In all modeling scenarios, we obtained major clusters of solutions (>98% of models) with a cluster precision of 21-25 Å (**Fig. S7A-D** and **Table S2**), each capable of supporting both globular and domain-swapped modes (**Fig. 4A**). An analysis of the overall fit to all DCX-DCX crosslinks shows a weak preference for the latter (**Fig. 4B**), but we cannot discriminate with high confidence based on utilization rates alone. However, an inspection of the centroid model for the main cluster of solutions shows a head-to-tail conformation with strong similarity to the domain-swapped structure (**Fig. 4C,D**). We note that no symmetry constraints were enforced during the modeling and it only is guided by crosslinking data. Taken together, there appears a preference for the open conformation through a domain swapping event in the CDC. Although dimerization through the globular domains remains possible, a conformational change induced by a lattice interaction could readily explain why dimerization in solution is not possible.

### DCX may not adopt a unique orientation on MTs

We next modeled the DCX-MT interaction with the complete set of crosslinks, to determine if the addition of crosslinks between DCX and MT (in particular) could orient the dimer on the MT lattice. We imposed a set of restrictions based on the findings described above. That is, for the full modeling exercise, we assumed that NDC binds to the MT lattice at the junctional binding site and DCX dimerizes on MT lattice specifically through CDC and C-tail regions. We carried over the degree of ambiguity in the nature of the dimerization event by modeling with both open and globular CDC structures and assessed the same four possible orientations for the dimer. For each of the resulting eight modeling exercises, one major cluster was obtained with a sampling precision of 14-20 Å, containing more than 99% of all the individual models (cluster precision of 26-31 Å) (**Fig. S8A-D** and Table S3). We could detect no preferred orientation for the DCX dimer on the lattice, based on crosslink usage. The distribution of crosslinks shows they accommodate all the orientations equally well (**Fig. 5A**). This dispersion suggests that an underlying heterogeneity in lattice engagement is possible.

**Fig. 5.**
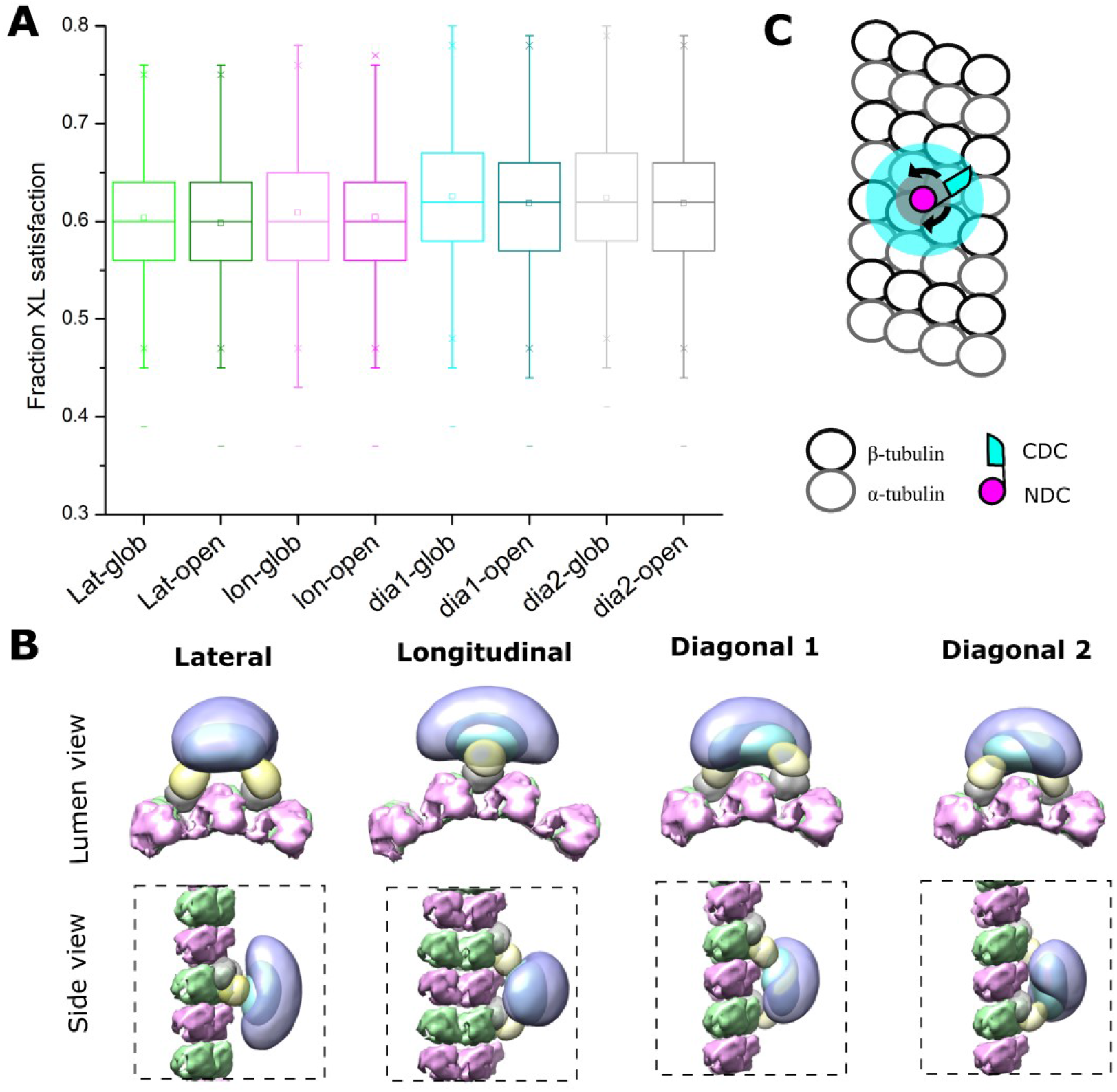
Positional evaluation of dimeric DCX on MT lattice. **A)** The fractional XL satisfaction for the main cluster of models generated for dimeric DCX-MT, employing all crosslinking restraints. The four different relative positions of fixed NDC on the MT lattice and the 2 CDC conformations were assessed. **B)** The density maps corresponding to the main cluster of the dimeric DCX-MT model, using all crosslinking restraints. The four different relative positions of fixed NDC on the MT lattice was assessed; α-Tubulin and β-Tubulin are shown in pink and green, respectively. NDC, linker, CDC and C-tail are shown as grey, lemon, cyan and light purple, respectively. **C)** The DCX-MT interaction model showing the flexibility of the DCX structure on the MT lattice.

Finally, to evaluate if the C-terminal tails engage the MT lattice in a preferred orientation, we inspected the density maps of the ensemble of models for each of the orientations (**Fig. 5B**). In all cases, regardless of the orientation, the tails are located distal to the lattice and adopt an averaged orientation perpendicular to the long axis of the dimer.

## Discussion

The function of DCX in MT regulation and brain development is dependent on the full-length protein and we propose that this function is strongly dependent upon its self-association. Our findings support a binding mode that presents DCX primarily as an intra-lattice dimer, oriented through NDC interactions at the junctional sites, and linked through the CDC domain. Based on the data-directed models that we generated in this study and the high fraction of crosslinking data that was satisfied, a lattice-induced dimerization event is the dominant state at the stoichiometry we explored. However, the association could adopt a variety of orientations on the stabilized MT lattice, consistent with an initial contact complex that involves just a monomer. That is, the flexibility of the interdomain region could support the precession of the monomer, allowing it to capture a suitably positioned additional monomer (**Fig. 5C**).

A primary binding mode involving CDC at the junctional sites is unlikely in the MT polymerization state that was studied here. Although we observed some crosslinks between the CDC to the MT lattice, most of these tend to locate the CDC away from the canonical site and may simply indicate couplings to the dimer as it transiently lays down on MT lattice. The interaction involving the C-tail domains is particularly intriguing and may indicate that self-association is more extensive than dimerization of the structured domains. An earlier observation highlighted strong binding cooperativity, revealing a Hill coefficient approaching 3 for DCX binding to 13-protofilament MTs (Bechstedt and Brouhard, 2012). This remarkable degree of cooperativity suggests that associations beyond dimeric are indeed possible and it likely involves the long C-tails. At the stoichiometry we explored, it is easy to imagine that these tails can associate across dimers to further stabilize the MT lattice (**Fig. 5B**).

The binding mode we describe in this study can unify interaction models that appear at first glance to be contradictory. Crystallographic evidence suggests that the CDC can dimerize under mildly denaturing conditions (Rufer et al., 2018) whereas cryo-EM data indicate that both NDC and CDC can bind to the lattice. A recent cryo-EM study by the Moores lab confirms that the CDC domain participates in MT nucleation at the junctional site but as the lattice grows, the sites become mostly occupied with NDC domains (Manka and Moores, 2020). The key to resolving this discrepancy is the conformational flexibility of the CDC. We propose a model where interactions between NDC and the MT lattice induce an opening of the CDC, leading to a binding event that favors the published crystallographic structure of the dimerized CDC (Rufer et al., 2018) (**Fig. 6**). The dimerization must be induced as it does not exist in solution without the MT interaction (Moores et al., 2006). The cryo-EM data seems to support this, as the conformational state of the CDC in the lattice-bound form is quite distinct from the X-ray structure of the free domain (Manka and Moores, 2020).

**Fig. 6.**
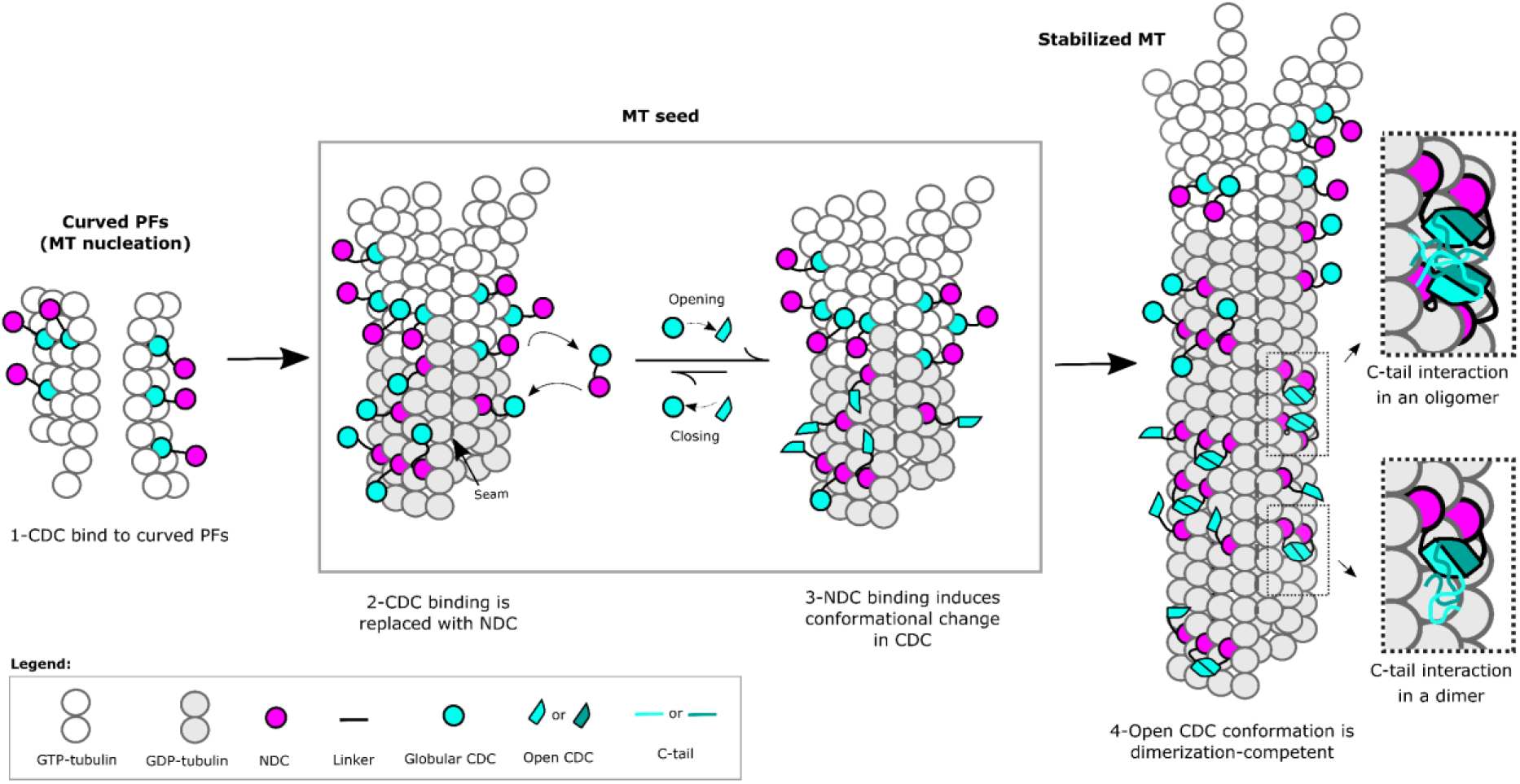
Mechanism of DCX-mediated MT nucleation and stabilization. DCX stabilizes early GTP-tubulin oligomers through the CDC domain, which is then replaced with the NDC domain at the canonical binding site as during full MT assembly. NDC binding triggers a conformational change in CDC domain, which facilitates DCX self-association and prevents CDC from rebinding. The inserts illustrate the role of C-tail domain in the formation of either inter-monomer interactions, inter-dimer interactions, or both.

Induced dimerization provides an explanation for the transition from an early CDC-MT interaction to an NDC-MT interaction (Manka and Moores, 2020) (**Fig. 6**). It was shown that CDC participates in nucleating MT growth when centrosomal γTURC complexes are not available, as is the case in the distal ends of migrating neurons. The deformable CDC domain creates junctional sites that may be somewhat different than the sites in the mature MT lattice because of the curved nature of the nucleated tubulin oligomers. As the lattice fully assembles and regular junctional sites are formed, the more rigid NDC domain can now bind. It is important to note that neither of the two structured domains bind to the lattice particularly strongly on their own, thus a transition to NDC binding could readily occur upon lattice maturation. The CDC interactions would diminish over time, provided that the structural features of the lattice prevent their rebinding, or a mechanism exists for their sequestering (or both). Our model suggests that sequestering is very likely to occur. Induced unfolding of CDC followed by its capture in a dimeric state would explain how DCX binds to the lattice so avidly and such an association would certainly prevent CDC from rebinding. At the same time, dimerization would stabilize the MT lattice and offer a compelling reason for favoring MTs with a well-defined number of protofilaments. That is, the more rigid NDC coupled with its larger footprint on the lattice (Manka and Moores, 2020) is better positioned than the CDC to set the angle of curvature, and dimerization increases the local concentrations of these domains on the lattice through avidity.

There are several functional observations that support the idea that dimerization helps define the overall MT architecture. A double NDC construct induces much more MT bundling than the wild-type (Manka and Moores, 2020), very likely through cross-lattice interactions, suggesting that dimerization minimizes these interactions. This is consistent with the conformation shown in **Fig. 5B**. A monomeric form may allow for bridging events leading to bundling, but our dimerization model would sterically hinder them. Dimerization is also supported through an inspection of patient mutations. These mutations cluster in both the NDC and CDC domains and lead to loss of MT binding and stabilization. It was suggested to be evidence that both domains interact with MTs directly (Taylor et al., 2000, Kim et al., 2003). Mutations in the NDC could certainly lead to reduced binding (Manka and Moores, 2020), but the mutations in the CDC region would more likely interfere with dimerization, reduce the avidity of DCX on the lattice and diminish MT stabilization overall.

Extended self-association involves the C-tail and additional observations support a role for this domain in regulating the MT architecture. Removing the C-tail reduces the preference for 13-protofilament MTs (Moores et al., 2004). Although its removal slightly increases DCX affinity to MTs, it also reduces the rate of DCX accumulation on the lattice (Moslehi et al., 2017). These findings suggest that the C-tail influences either the span or the stability of the dimerization event. The depth of crosslinking we observed in this region suggests an induced structure, perhaps a coiled-coil as observed with other dimerized MAPs (Slep et al., 2005, Kon et al., 2009). However, DeepCoil (Ludwiczak et al., 2019) shows a very low probability of such a domain. A more extensive multi-DCX network backed by a dynamically interacting set of C-tails would explain the high cooperativity of binding and help stabilize DCX on the lattice (see inserts in **Fig 6**). It is possible that the C-tails may influence the NDC-MT interaction in a more direct fashion but either way, the self-association of the C-tails affect the properties of DCX on the lattice. This effect can be regulated by C-tail phosphorylations (Tanaka et al., 2004, Graham et al., 2004, Shmueli et al., 2006). For example, phosphorylation of S297 by Cdk5 reduces DCX affinity to MTs in cultured neurons and any mutations (phosphomimics or dephosphomimics) similarly reduce affinity and alter neuronal migration patterns (Tanaka et al., 2004). Conversely, spinophilin-induced dephosphorylation of S297 by PP1 causes an increase MT bundling in the axonal shaft of neurons (Bielas et al., 2007). Collectively, these observations indicate that higher order self-association and its regulation are critical to DCX function on the MT lattice.

At this point, it is unclear if the precise orientation of the dimer on the MT lattice is important or not. DCX recognizes a specific tubulin spacing in the lattice, favoring longitudinal curvatures that result in shorter inter-dimer distances (Ettinger et al., 2016). It suggests that a longitudinal orientation is possible, but this particular property may be a function of direct CDC binding at curvatures (in monomeric form, **Fig. 6**), which is consistent with a role in sensing MT plus-end geometries (Bechstedt et al., 2014, Manka and Moores, 2020). Missense mutations in the CDC region (and not NDC) disrupt the longitudinal curvature detection properties of DCX, but this could influence dimerization as well. Our integrative modeling is not sufficiently precise to identify a dominant orientation (if one truly exists), but a lateral interaction seems the most likely given the orientation of the NDC-CDC linker in cryo-EM models (Manka and Moores, 2020). A dimer with an ability to engage the lattice in a lateral manner could reinforce a preference for a 13-protofilament lattice, if the “bite length” of the dimer is complementary to protofilament spacing.

Finally, the success of this integrative modeling technique relies heavily upon an improved crosslinking methodology that samples the equilibrated state more faithfully (Ziemianowicz et al., 2018). The presence of nominally disordered regions in MAPs like DCX renders proteins prone to “kinetic trapping” on the lattice, where conventional long-lived crosslinking reagents can trap interactions in non-native states. We conducted a modeling analysis of the DCX-MT interaction using typical DSS and EDC crosslinkers and while the data are abundant (**Fig. S9A,B**), the outcome of the integrative modeling using conventional crosslinking chemistries was poor, both in terms of the percent crosslink satisfaction (<50%) and the location of the primary binding site (**Fig. S9C-F**). The full modeling exercise with these data provided no interpretable results. While a small number of studies have applied crosslinking to MAP-MT interactions (Legal et al., 2016, Zelter et al., 2015, Abad et al., 2014, Kadavath et al., 2015), many more could be profiled with an improved crosslinking method such as we illustrate here.

## Methods

### Expression and purification of DCX

The production method for DCX was described previously (Bechstedt and Brouhard, 2012). Briefly, the plasmid for human DCX (sp|O43602|DCX_HUMAN/1-365) includes an N-terminal polyhistidine (6His) tag and a C-terminus StrepTag II. DSB128 (DCX-pHAT-HUS strain in BL21) was grown in 100 mL 2YT supplemented with 100 µg/mL ampicillin and 20 µg/mL gentamycin at 32°C overnight to OD600 ∼1-3. Cells were resuspended in fresh media and grown at 37°C to mid-log phase (OD600 ∼ 0.8 – 1.0). DCX expression was induced by addition of IPTG and incubated for 16 h at 18°C. Cells were harvested by centrifugation and stored at -80°C until use.

For the expression of isotopically labeled DCX, DSB126 (DCX-pHAT-HUS in Arctic Express) was grown as above to OD_600_ ∼1-3. The rich 2YT media was removed and cells resuspended in 1L of freshly prepared ^15^N-M9 minimal media (Lima et al., 2018), with 100 μg/mL ampicillin and 20 µg/mL gentamycin. The cells were then grown at 37°C to mid-log phase (OD_600_ ∼ 0.8), the temperature reduced to 16°C for 1 h, then induced with 1 mM IPTG overnight. Cells were harvested by centrifugation and stored at -80°C until use.

For protein purification, for either state, DCX-expressing cells were thawed and resuspended in cold lysis buffer (50 mM Tris-HCl pH 8, 300 mM NaCl, 10 mM imidazole, 10% glycerol, cOmplete protease inhibitor, 1 mM PMSF) and lysed by sonication (QSonica, Newtown, USA) on ice. After centrifugation, the supernatant was loaded onto a HisTrap column (5 mL, cat. No. 17-5248-01, GE Healthcare) pre-equilibrated with lysis buffer. The column was washed with His-Buffer A (50 mM Tris-HCl pH 8, 300 mM NaCl, 10 mM imidazole, 1 mM PMSF) a protein eluted with a gradient of 0-100% His-Buffer B (50 mM Tris-HCl pH 8, 300 mM NaCl, 250 mM imidazole, 1 mM PMSF). Fractions were collected and analyzed by Western blot, probed with anti-His (Applied Biological Materials Inc, Richmond, Canada) and Streptavidin-HRP (ThermoFisher Scientific). DCX-containing fractions were dialyzed (MWCO 6-8kDa) in Strep-Buffer A (100 mM Tris-HCl pH 8, 150 mM NaCl, 1 mM EDTA) at 4^°^C overnight. The dialyzed DCX sample was centrifuged to remove any precipitates before loading onto a Strep-Trap (1 mL, cat. No. 29-0486-53, GE Healthcare) column. After washing with Strep-Buffer A, DCX was eluted with a gradient of 0-100% Strep-Buffer B (100 mM Tris-HCl pH 8, 150 mM NaCl, 1 mM EDTA, 2.5 mM D-Desthiobiotin, 10% glycerol (VWR Life Science)). Fractions were analyzed by Western blot as above and DCX-containing fractions were dialyzed in BRB80 (80 mM PIPES, 1 mM EGTA,1 mM MgCl_2_, pH 6.8) and supplemented with 1 mM GTP (Enzo Life Sciences, Farmingdale, USA). After centrifugation, the supernatant was concentrated on an Amicon MWCO 10 kDa unit (Millipore). The concentration of DCX was determined by Nanodrop (ThermoFisher)

### Fluorescence microscopy

Porcine Tubulin at 1:5 ratio (rhodamine-labeled: unlabeled) (cat. no. T240 and TL590M-A respectively, Cytoskeleton, Inc., Denver, USA) was reconstituted to 4.0 mg/mL in cold polymerization buffer, BRB80. Centrifugation was performed at 16,800 g for 10 minutes at 4°C to pellet any aggregates. The supernatant was diluted in warm BRB80 to 10 µM (1.0 mg/ml) and supplemented with 10 µM DCX and incubated at 37°C for 90 minutes. The resulting DCX-MT constructs were diluted 100 times in warm resuspension buffer (BRB80, 10 μM Docetaxel) immediately before fluorescence microscopy. Diluted sample was placed on a glass slide and covered with cover glass (glass slides were acid-etched overnight before use and rinsed thoroughly). MTs were imaged using an AxioObserver epi-fluorescence microscope with an oil immersion objective (EC Plan Apo 100x/1.3, Carl Zeiss MicroImaging GmbH, Jena, Germany) equipped with CCD camera Zeiss AxioCam MRm Rev. 3 FireWire. Images were taken using 100% light source intensity and 4 s exposure time. Microscopy images were processed to adjust for contrast using Zen Blue software (version 2.3). MT length measurement were performed using standard measurement tools in ImageJ (Schindelin et al., 2012).

### Turbidity assay

Tubulin (10 µM) and different concentrations of DCX (0 – 20 µM) were premixed in BRB80 + GTP (1 mM) on ice and transferred to a 396-well plate. The plate was transferred to the temperature controlled (37°C) spectrophotometer (Molecular Devices FilterMax F5 spectrophotometer equipped with a 340 nm filter). Turbidity measurements were taken every 13 s for over 70 minutes. Data were normalized to a preliminary time point and the pathlength was corrected to 1 cm.

### Crosslinking

Porcine Tubulin was reconstituted to 4.0 mg/mL in cold BRB80 polymerization buffer supplemented with 1 mM GTP. Centrifugation was performed at 16,800 g for 10 minutes at 4°C to pellet any aggregates. The supernatant was diluted to 1.0 mg/mL (10 µM) in the presence of 10 µM DCX and incubated at 37°C for 90 minutes to induce polymerization. Succinimidyl 6-(4, 4’-azipentanamido)hexanoate (LC-SDA; Thermo Scientific) crosslinking was performed by adding the reagent to final 1mM concentration. The sample was incubated for 10 minutes at 37°C, followed by 5 sec of photolysis at 355 nm (50 x 100 mJ laser pulse of a 10 ns pulse-width using an Nd:YAG laser, YG 980; Quantel, Les Ulis, France) (Ziemianowicz et al., 2018). DSS crosslinking was performed using 1mM crosslinker concentration followed by 30-minute incubation at 37°C. EDC crosslinking was performed with the same crosslinker concentration and incubation time, in the presence of 2mM Sulfo-NHS. All chemicals were purchased from Sigma Aldrich, unless mentioned otherwise.

### Mass Spectrometry

Crosslinked DCX-MT was separated from free protein by centrifugation. The supernatant was removed, and the pellet washed once with warm BRB80 (37°C) and then dissolved in 50 mM ammonium bicarbonate solution to a final protein concentration of 1 mg/ml. Cysteines were reduced by adding DTT to a final concentration of 10 mM with heating to 90°C for 10 min, and then alkylation by addition of chloroacetamide to a final concentration of 50 mM at 37°C for 30 min. Trypsin (Thermo Scientific) was added to an enzyme-to-substrate ratio of 1:50 and incubated overnight at 37°C with nutation (150 rpm). Digestion was quenched by adding formic acid to a final concentration of 2%. Samples were aliquoted and either lyophilized for size-exclusion chromatography (SEC) or desalted using ZipTip C18 pipette tips (Merck Millipore Ltd. Ireland) for LC-MS/MS analysis. For SEC, samples were reconstituted in SEC buffer consisting of 30% acetonitrile (LC-MS grade, Thermo Scientific) and 0.1% FA (Fluka) and crosslinked peptides were enriched by separation on a Superdex Peptide PC 3.2/30 column (GE Healthcare), using an Agilent 1100 chromatography system at 95 μL/minute with fraction collection. Fractions were lyophilized. SEC fractions and un-enriched samples were reconstituted in 0.1 % formic acid and based on the UV absorbance trace, approximately 1 pmol injected onto a Acclaim PepMap 100 guard column (75 μm×2 cm C18, 3 μm particles, 100 Å; Thermo Scientific) and separated on a 50 cm PepMap RSLC C18 (75 μm×50 cm, 2 μm particles, 100 Å; Thermo Scientific) coupled to an Orbitrap Fusion Lumos (Thermo Scientific). Peptides were separated with a gradient consisting of 56 minutes at 5-28% B, 17 minutes at 28-40% B, followed by 10 minutes 40-95% B and regenerated at 95% B for 10 minutes. Mobile phase A was 0.1% FA in water, and mobile phase B was 0.1% FA in 80% ACN. The flow rate was 300 nL/minute. The mass spectrometer was operated in positive ion mode, in OT/OT mode with MS resolution set at 120,000 (350–1300 m/z) and MS2 resolution at 15000. The max injection time was set at 50 ms and 100 ms for MS1 and MS2 scans, respectively. The AGC target was set at 400,000 and 100,000 for MS1 and MS2, respectively. Higher energy collisional dissociation (HCD) was used to generate MS2 spectra, for charge states 4+ and higher. A normalized collision energy of 32% was used with an isolation width of 1.2 m/z.

### DCX dimerization analysis

^14^N-DCX (light) and ^15^N-DCX (heavy) was mixed in 1:1 ratio and then applied to tubulin as above. To explore dimerization and/or aggregation, the light-heavy mixture was processed using a few different approaches: 1) mixture incubated at 10 µM for 2h at 37°C with tubulin, 2) mixture incubated at 10 µM for 2h at 37°C without tubulin, 3) mixture diluted to 2 µM followed by concentration to ≥10 µM and incubation at 37°C for 2h without tubulin, 4) mixture denatured in 3M Guanidyl-HCl in PBS overnight followed by buffer exchange into BRB80 and concentration to ≥10 µM and incubation at 37°C for 2h without tubulin. A negative control sample was also prepared by crosslinking heavy and light DCX separately and mixing them immediately prior to LC-MS/MS analysis. All samples were crosslinked by LC-SDA and processed as above, using only the non-enriched samples.

### XL-MS data analysis

Raw LC-MS/MS data were imported into CRIMP v2 (the crosslinking plugin in the Mass Spec Studio, www.msstudio.ca) (Sarpe et al., 2016) along with the Fasta files of the Tubulin isoforms identified previously (TB-α-1A, TB-α-1B, TB-α-1C, TB-α-1D, TB-α-4A, and TB-β, TB-β-2B, TB-β-4A, TB-β-4B, TB-β-3, TB-β-5) (Rafiei and Schriemer, 2019). The minor impurities in the recombinantly purified DCX did not have an impact on crosslinking results. For crosslinked peptide searching, methionine oxidation and carbamidomethylation of cysteines were selected as variable and fixed modifications, respectively. Crosslinked peptides were searched using the following parameters: MS accuracy, 5 ppm; MS/MS accuracy, 10 ppm; E-threshold, 70; Enzyme, Trypsin (K/R only); m/z range: 350 and 1300; Peptide length range: 3-50 residues. All crosslinking results filtered at an estimated 1% FDR level were manually validated. Monoisotopic ion identification in MS1 was confirmed in all hits and only crosslinked peptides with a good fit of the measured isotopic envelope to the theoretical isotopic pattern were accepted. Crosslinking data were exported as CSV files and data from all SEC fractions combined. For isotopically labeled proteins, CRIMP was customized to search all heavy-light peptide combinations.

### DCX-MT integrative structure modeling

We used integrative structure modeling (ISM) to assess how our generated crosslinking data, along with the previously computed EM map, prior structural information and physical principles, could model the interaction between the domains of DCX and the microtubule lattice (SI Methods for details). Briefly, data and information were transformed into spatial restraints used to construct a scoring function and then we searched for model configurations that satisfy all restraints. To assess whether NDC or CDC binds to MT, we performed ISM using the MT lattice and either CDC or NDC along with intermolecular crosslinks between each domain and MT. The convergence of each of the DCX domains to the observed binding site in the EM map using only the crosslinking data was analyzed. In addition, the satisfaction of the crosslinking restraint was computed while enforcing a restraint using the EM density of the domain of DCX bound to MT to guide the domain to this observed site, with the better models identified by those that better satisfy the crosslinks.

We assessed the nature of the DCX dimerization event by performing ISM using two DCX molecules and a 3-4 repeat (lateral) and 3-4 repeat (longitudinal) MT domain constructed using tubulin and symmetry operations. The position of NDC (NDC structure of PDB: 4ATU (Liu et al., 2012)) was fixed on the MT lattice in the site observed by EM. CDC was modeled in both the open (domain-swappable, PDB 6FNZ (Rufer et al., 2018)) and closed (globular structure, PDB: 5IP4 (Burger et al., 2016a) forms. Models were scored via the set of all DCX-DCX crosslinks and models were assessed using crosslink satisfaction. Finally, we evaluated the ability of DCX dimers to be formed using four spatial arrangements of DCX dimers on the MT lattice. Two copies of NDC were fixed in adjacent binding sites Lateral, Longitudinal, Diagonal-1 and Diagonal-2 (**Fig. S10**). Models were generated using restraints based on all cross-linking data (DCX-DCX and DCX-MT).

Integrative structure modeling protocols were built using the Python Modeling Interface (PMI) package of the open-source Integrative Modeling Platform (IMP) package (Webb et al., 2018) version 2.10 (https://integrativemodeling.org). Specifically, integrative structure modeling proceeds through four stages: 1) gathering data, 2) representing subunits and translating data into spatial restraints, 3) sampling configurations of the system representations to produce an ensemble of models that satisfies the restraints, and 4) analyzing and validations the ensemble structures (**Fig. S2**).

#### Stage 1: Gathering data

The 124 unique crosslinks identified between DCX and α−β tubulin, along with 74 unique inter- and intra-DCX crosslinks identified in this study informed the spatial proximities of the CDC and NDC with respect to microtubule lattice and each other. An EM map containing MT and DCX (EMDB 2095 (Liu et al., 2012)) at 8.0 Å resolution was used to inform the binding mode of DCX with respect to the MT lattice. The map was prepared as follows. First, the cryo-EM structure for MT polymerized in the presence of DCX (6EVZ (Manka and Moores, 2018)), was fitted into the EM map using UCSF Chimera tools. Second, the EM volumes corresponding to the MT structure were erased using Erase tool in Chimera, leaving only the DCX density. The representation of the microtubule lattice was derived from the high-resolution structure built based on the cryo-EM structure for MT polymerized in the presence of DCX (PDB: 6EVZ (Manka and Moores, 2018)). The representation of the NDC globular domain was informed by the PDB structure 4ATU (Liu et al., 2012) and the representation of the globular domain of CDC informed by either the crystal structures of the globular (PDB 5IP4 (Burger et al., 2016b)) and domain-swapped (PDB 6FNZ (Rufer et al., 2018)) configurations.

#### Stage 2: System representation and translation of data into spatial restraints

The information above was used to define the representation of the system, the scoring function that guides the search for configurations that satisfy the information, filtering of models and validation of the final ensembles. One representation of the MT lattice was constructed based on the coordinates of the PDB structure (6EVZ (Manka and Moores, 2018)), with components simultaneously represented as beads of a single residue each and up-to-10 residues-per-bead. An additional 4-dimer width MT lattice representation was made using the same PBD structure (6EVZ) of MT lattice and symmetry replication tools in UCSF Chimera (Kim et al., 2003).

Residues of NDC (51-140) and CDC (174-251 for globular and 174-253 for open form) contained in the crystal structure (structured domains) were represented simultaneously as beads of up-to-10 residues-per-bead. Residues not contained in the crystal structures (terminal tails and the linker region between NDC and CDC were modeled solely as beads of up-to-10 residues each. Both the NDC and CDC structured domains were constrained as rigid bodies: the intermolecular distances between these beads were kept fixed. All other beads were unconstrained. The N-terminal (1-50) and C-terminal (331-360) tails of doublecortin were not modeled, as these regions were less represented in inter-and/or intra-DCX XL sites. For different modeling protocols, either one of the NDC/CDC was modeled, or two copies of DCX modeled.

The information was then encoded into spatial restraints computable on the system representation. Our scoring function consisted of the sum of four component restraints. 1) The *chemical cross-link restraint* utilized the identified chemical cross-links to construct a Bayesian scoring function (Gutierrez et al., 2020) that restrained distances between cross-linked residues. Restraint distances of 30Å for LC-SDA and DSS, and 25Å for EDC were used. The ambiguity of crosslink sites due to presence of multiple copies of the same protein was considered. In these cases, the joint probability of satisfying the restraint over all possible combinations is computed (Molnar et al., 2014). 2) The *EM restraint* utilized a Bayesian scoring function based on the cross-correlation of the overlap of the model and experimental density. Densities of the model components and experimental data were approximated using a Gaussian Mixture Model (GMM) (Bonomi et al., 2019). 3) The *excluded volume restraint* prevents parts of the system from occupying the same space. This restraint is applied to the low-resolution representation of the system (10 residues per bead). 4) The *sequence connectivity restraint* was used to restraint chemical components known to be covalently linked. The restraint was applied as a harmonic upper-distance bound on the distance between adjoining beads in sequence. The center of the harmonic was defined as twice the sum of the radii of the two beads, the radii computed from the excluded volume of the bead.

#### Stage 3: Configurational sampling

The search for model configurations that satisfy the restraints used Gibbs sampling, based on the Metropolis Monte-Carlo algorithm and accelerated via replica exchange. Initial positions of system components were randomized with exception of MT being fixed for all cases, and NDCs being fixed for dimeric-DCX modeling scenarios. The set of Monte-Carlo moves consisted of random rotation and translation of rigid bodies (up to 4.0 Å and 0.5 radians, respectively) and random translations of the individual beads not in rigid bodies of up to 4 Å. Model configurations were saved every 10 steps. For each modeling protocol, 8-10 independent sampling runs were initiated using 20-40 replicas and run for 100,000-500,000 steps each, resulting in 100,000-500,000 models for each protocol.

#### Stage 4: Analyzing and validating model ensembles

The resulting ensemble of model configurations were analyzed to estimate the structural precision, ensure appropriate consistency with the input data and suggest more informative future experiments. The models were first filtered for those that satisfy the input data. After filtering, we used analysis and validation protocols published previously [25]. Briefly, analysis began by assessing the thoroughness of structural sampling and computation of the sampling precision, the RMSD threshold at which clustering produces indistinguishable results for two independent sets of computed models (for example, the output of runs 1-5 and 6-10), using protocols previously described (Viswanath et al., 2017). Upon computation of the sampling precision, the entire set of models was clustered at this precision or higher to ensure that resulting model clusters reflect the uncertainty in sampling in input information. The centroid models of clusters were then filtered to remove the models where NDC/CDC is located on the MT edges (where another tubulin monomer will fit in a full MT structure). These model clusters were further assessed by computing their precision (structural variability – average RMSD to the centroid model) and quantifying their fit to the input information.

These final models were validated by ensuring that the models satisfied the cross-linking data used to compute it. Crosslink satisfaction was also used to determine which modeling configurations were more plausible than others. The fraction crosslink satisfaction was computed both for each individual model configuration in a cluster and the entire cluster. A crosslink was deemed to be satisfied if the crosslink distance in an individual model configuration was less than 35 Å for LC-SDA and DSS and 30 Å for EDC. A crosslink was deemed to be satisfied in a model cluster (called cluster crosslink satisfaction in **Table S2**) if at least one configuration in the cluster has a crosslink distance less than the distance thresholds described above. Also, RMSD calculation was performed between each individual model present in the clusters and the NDC component of PDB structure 4ATU. RMSD calculation for CDC-MT models was performed using an aligned globular CDC structure (PDB 5IP4) into the NDC component of cryo-EM based structure (PDB 4ATU). The full details of sampling precision, clustering threshold, cluster’s population, etc. for all modeling scenarios are presented in **Table S2**.

## Data availability

The DCX-MT integrative models, including final structures, modeling details, and input experimental data, were deposited into the PDB-dev repository for integrative models (www.pdb-dev.com) accession number PDBDEV_00000071, PDBDEV_00000072, PDBDEV_00000073 and PDBDEV_00000074. All LC-MS/MS data generated to support the findings of this study have been deposited with the ProteomeXchange Consortium via the PRIDE (Perez-Riverol et al., 2019) partner repository with the dataset identifier PXD023950. Reviewer account details:

**Username**

**Password**

MiR4m61y

## Acknowledgments

We thank Dr. Laurent Brechenmacher for consultation in LC-MS/MS method development. This work was supported by CANARIE (project RS-326) and NSERC (RGPIN-2017-04879).

## SUPPLEMENTARY FOR

**Fig. S1.**
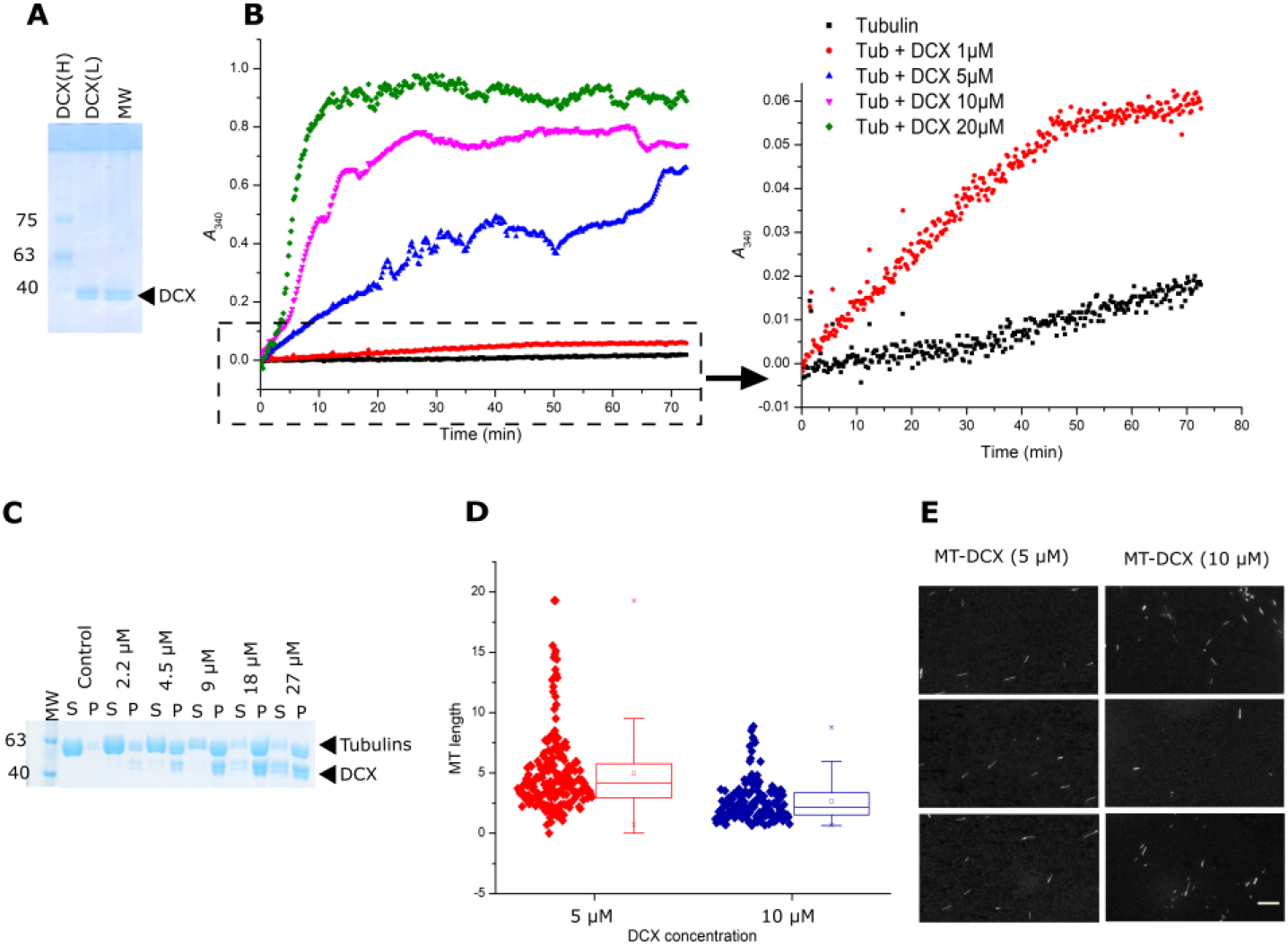
The evaluation of purity and activity of in house purified Doublecortin. **A)** The purity of DCX-Light and DCX-Heavy isotopically labeled sample was evaluated on 8% SDS-PAGE gel. **B)** Turbidity measurement at 340 nm versus time for Tubulin (10 μM) with different concentrations of DCX (1, 5, 10 and 20 μM) in the presence of 1mM GTP. Samples containing Tubulin with even low DCX concentrations produced a significant augmentation in scattering, confirming the formation of microtubules (MTs). The right graph is the zoomed-in view of the negative control containing Tubulin (10 μM) + GTP (1 mM) in black and Tubulin + DCX (1 μM) + GTP (1 mM) in red. **C)** MT polymerization in the presence of different concentrations of DCX (2.5-30 μM). The tested Tubulin concentration was 10 μM and GTP concentration was 1 mM. The control lane is devoid of DCX. **D)** The length distribution of MTs polymerized in the presence of 5 and 10 μM DCX concentrations. There is a significant difference between the two populations at p= 0.001. The horizontal line indicates the median value, the box indicates the 25th and 75th percentiles (N∼120). **E)** Fluorescence microscopy evaluation of MTs polymerized in the presence of DCX at 5 and 10 μM concentrations. No bundling was observed. scale bar = 10 µm.

**Fig. S2.**
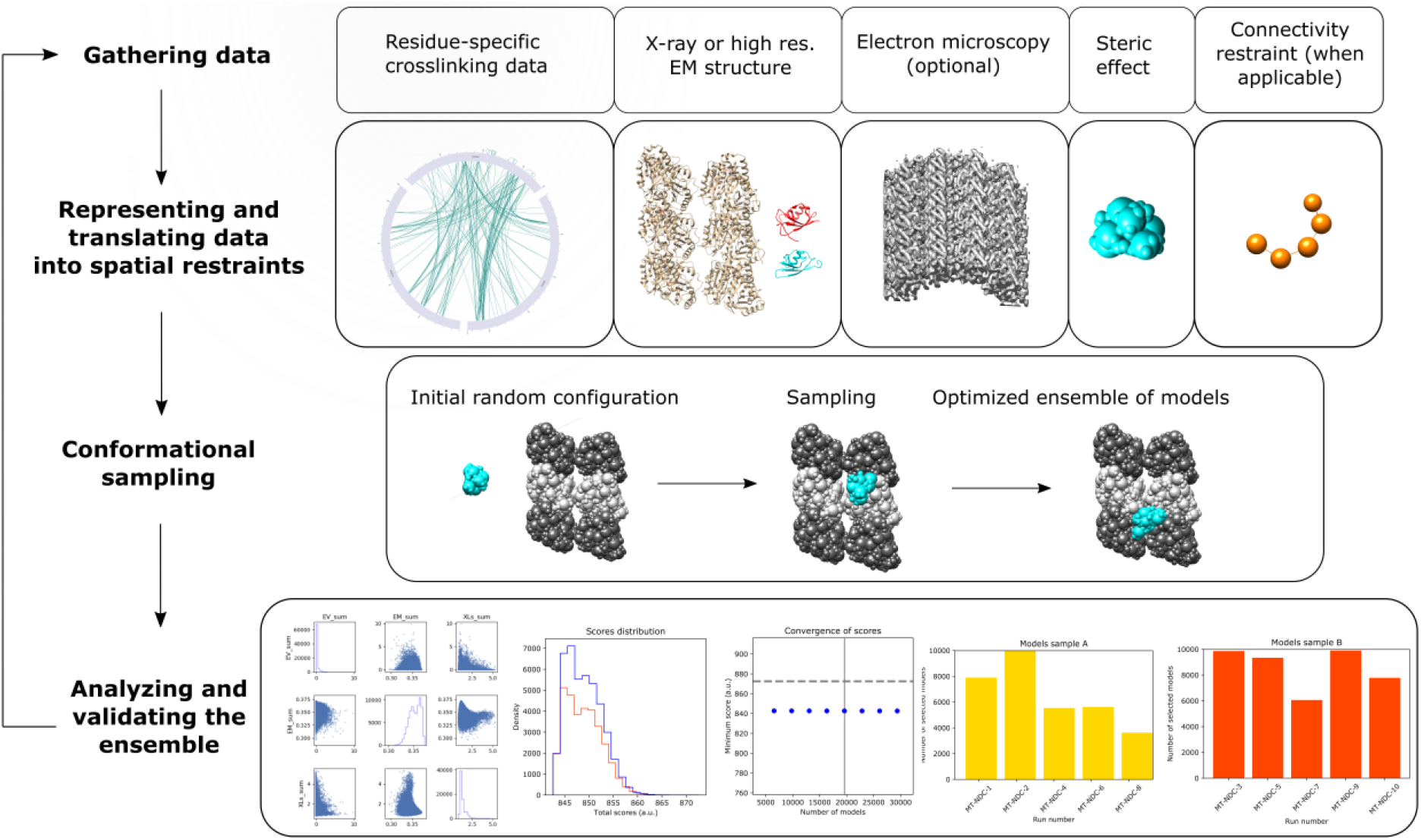
Integrative structure determination of MT-DCX, using four stage workflow: 1) Data gathering, including chemical crosslinking, atomic structures (MT structure, PDB 6EVZ [2]; NDC structure, DCX component of PDB 4ATU [1], CDC structure, PDB 5IP4 [3]), and cryo-EM map EMD 2095 [1]. 2) Representation of subunits and translation of the data into spatial restraints. 3) Conformational sampling to obtain an ensemble of structures needed to satisfy the input data. 4) Clustering the sample models into distinct groups of structures, followed by the analysis in terms of accuracy and precision.

**Fig. S3-A.**
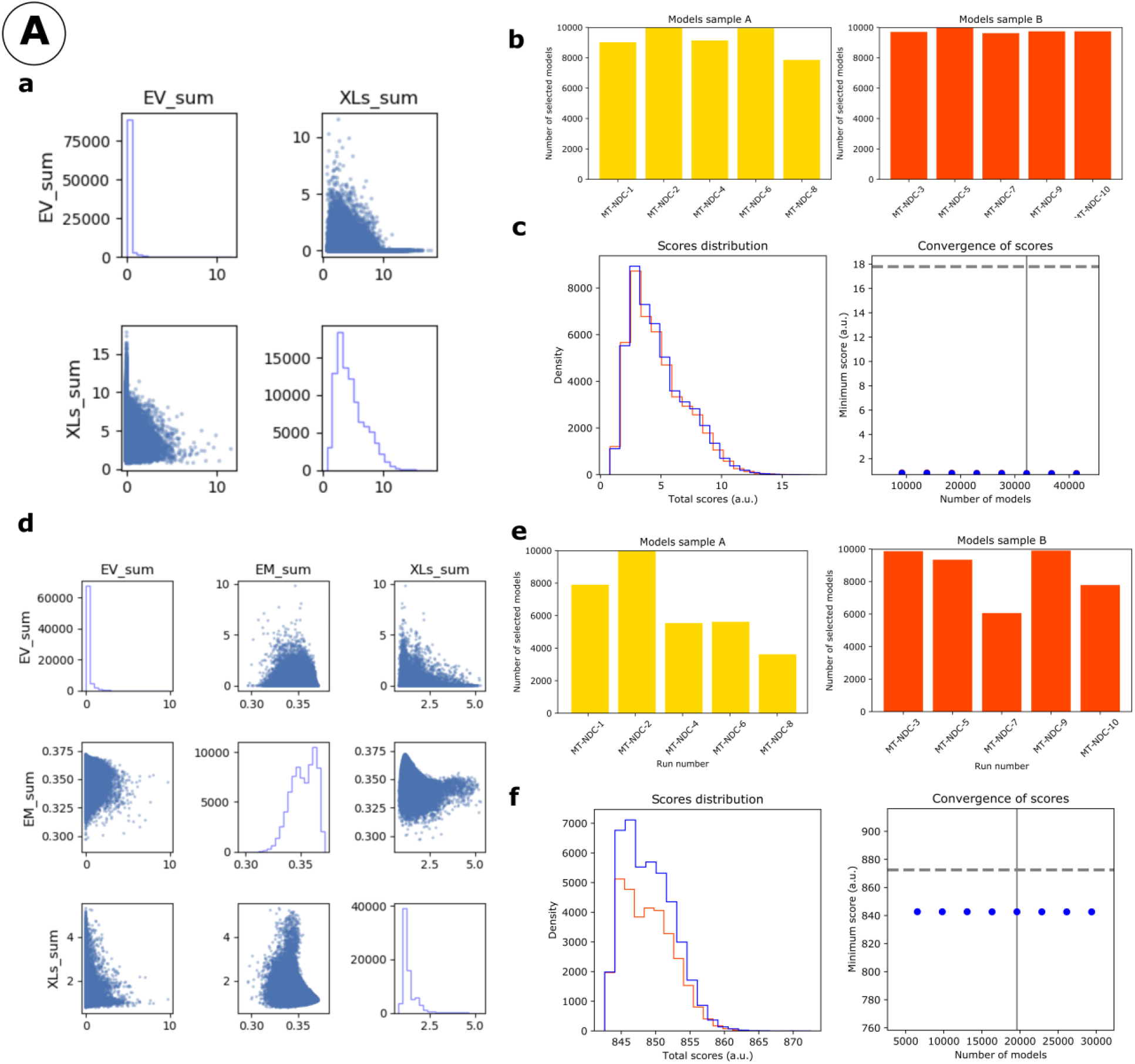
IMP analysis output for MT-NDC. **a)** The distribution of scores for the models computed for MT-NDC, using only crosslinking data (500,000 models). **b)** The number of models randomly selected for model samples A (yellow) and sample B (red), from 10 different modeling runs. **c)** The score convergence for models in samples A and B. The nonparametric Kolmogorov–Smirnov two-sample test (two-sided) indicates that the difference between the two score distributions is insignificant, the magnitude of the difference is small, as demonstrated by the Kolmogorov–Smirnov two-sample test statistic, D, of 0.006. Thus, the two score distributions are effectively equal. **d)** The distribution of scores for the models computed for MT-NDC, using both crosslinking data and EM map (500,000 models). **e)** The number of models randomly selected for model samples A (yellow) and B (red), obtained from 10 different modeling runs. **f)** The score convergence for the models in samples A and B. The nonparametric Kolmogorov– Smirnov two-sample test (two-sided) indicates that the difference between the two score distributions is insignificant, the magnitude of the difference is small, as demonstrated by the Kolmogorov–Smirnov two-sample test statistic, D, of 0.031. Thus, the two score distributions are effectively equal.

**Fig. S3-B.**
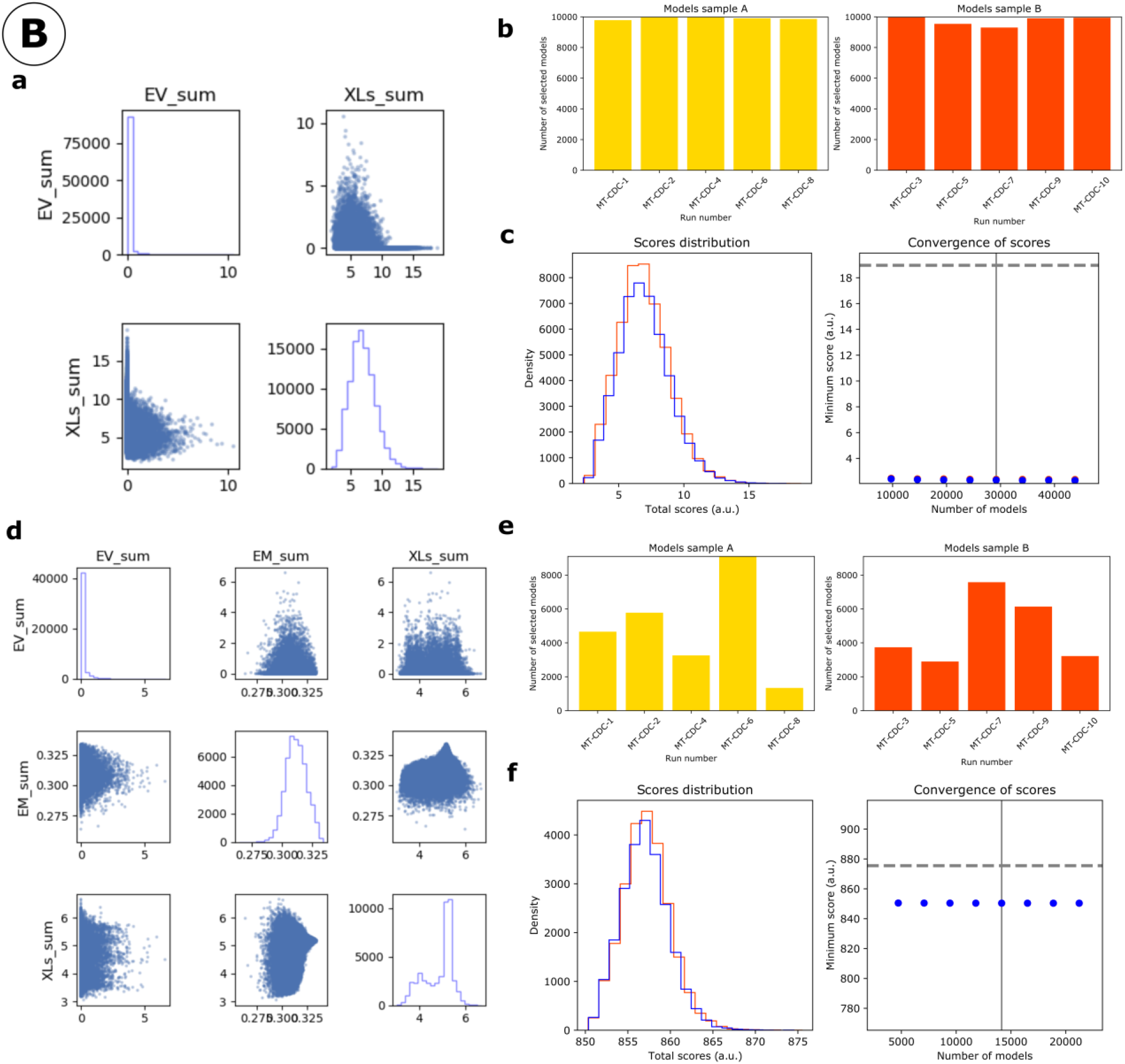
IMP analysis output for MT-CDC. **a)** The distribution of scores for the models computed for MT-CDC, using only crosslinking data (500,000 models). **b)** The number of models randomly selected for model samples A (yellow) and B (red), from 10 different modeling runs. **c)** The score convergence for models in samples A and B. The nonparametric Kolmogorov–Smirnov two-sample test (two-sided) indicates that the difference between the two score distributions is insignificant, the magnitude of the difference is small, as demonstrated by the Kolmogorov–Smirnov two-sample test statistic, D, of 0.010. Thus, the two score distributions are effectively equal. **d)** The distribution of scores for the models computed for MT-CDC, using both crosslinking data and EM map (500,000 models). **e)** The number of models randomly selected for model samples A (yellow) and B (red), from 10 different modeling runs. **f)** The score convergence for models in samples A and B. The nonparametric Kolmogorov–Smirnov two-sample test (two-sided) indicates that the difference between the two score distributions is insignificant, the magnitude of the difference is small, as demonstrated by the Kolmogorov–Smirnov two-sample test statistic, D, of 0.034. Thus, the two score distributions are effectively equal.

**Fig. S4.**
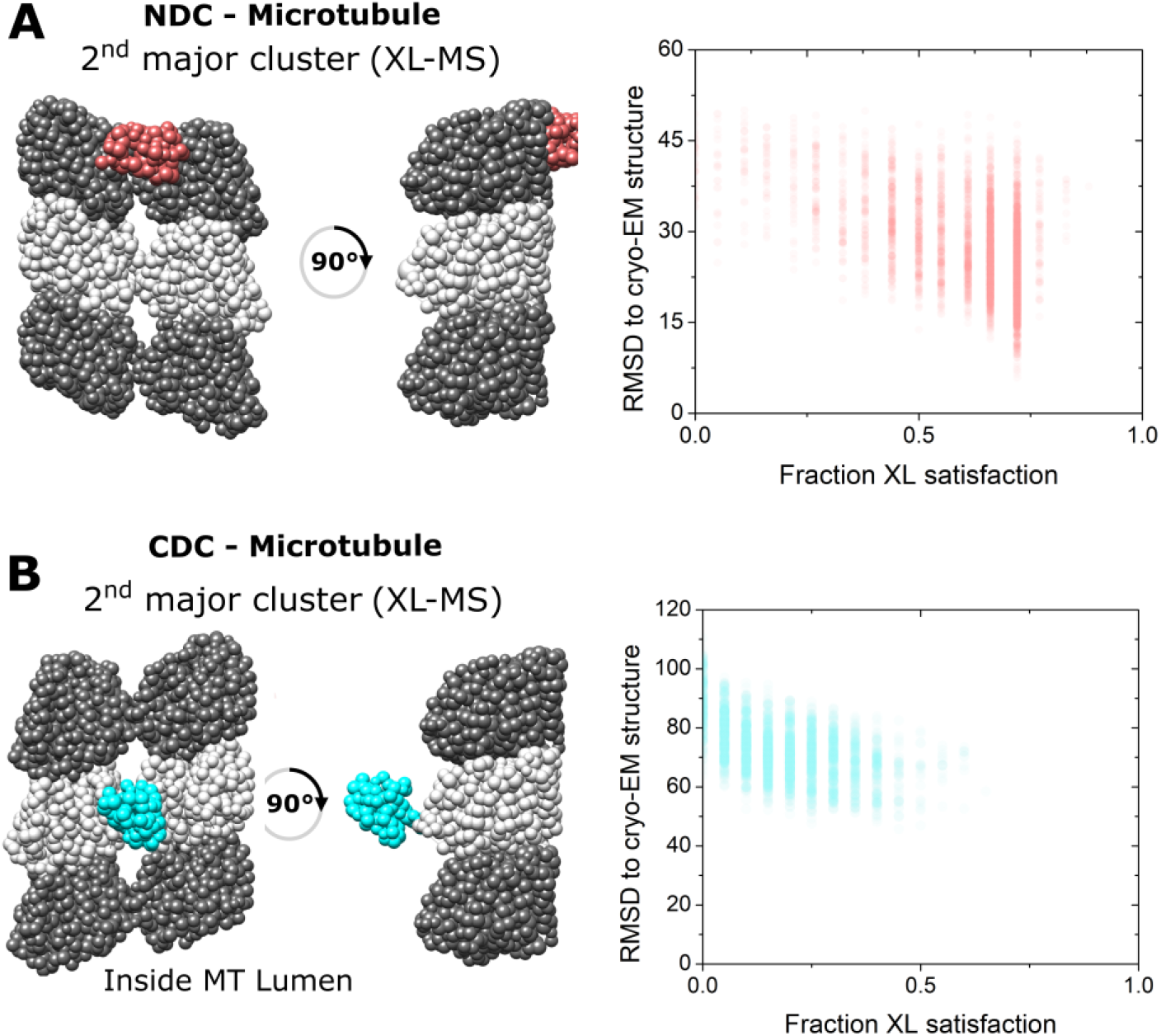
Integrative structural modeling for NDC-MT and CDC-MT (2^nd^ major clusters). The centroid structure for the 2^nd^ major cluster of models produced by IMP for NDC-MT (A) and CDC-MT (B) guided exclusively by XL-MS restrains; α−Tubulin and β−Tubulin are shown as light and dark grey, respectively. NDC is shown in red, and CDC is shown in cyan. The fraction crosslink satisfaction (defined as <35 Å) versus RMSD to canonical binding site (PDB 4ATU) for all models present in the main structural cluster is presented for each modeling scenarios.

**Fig. S5.**
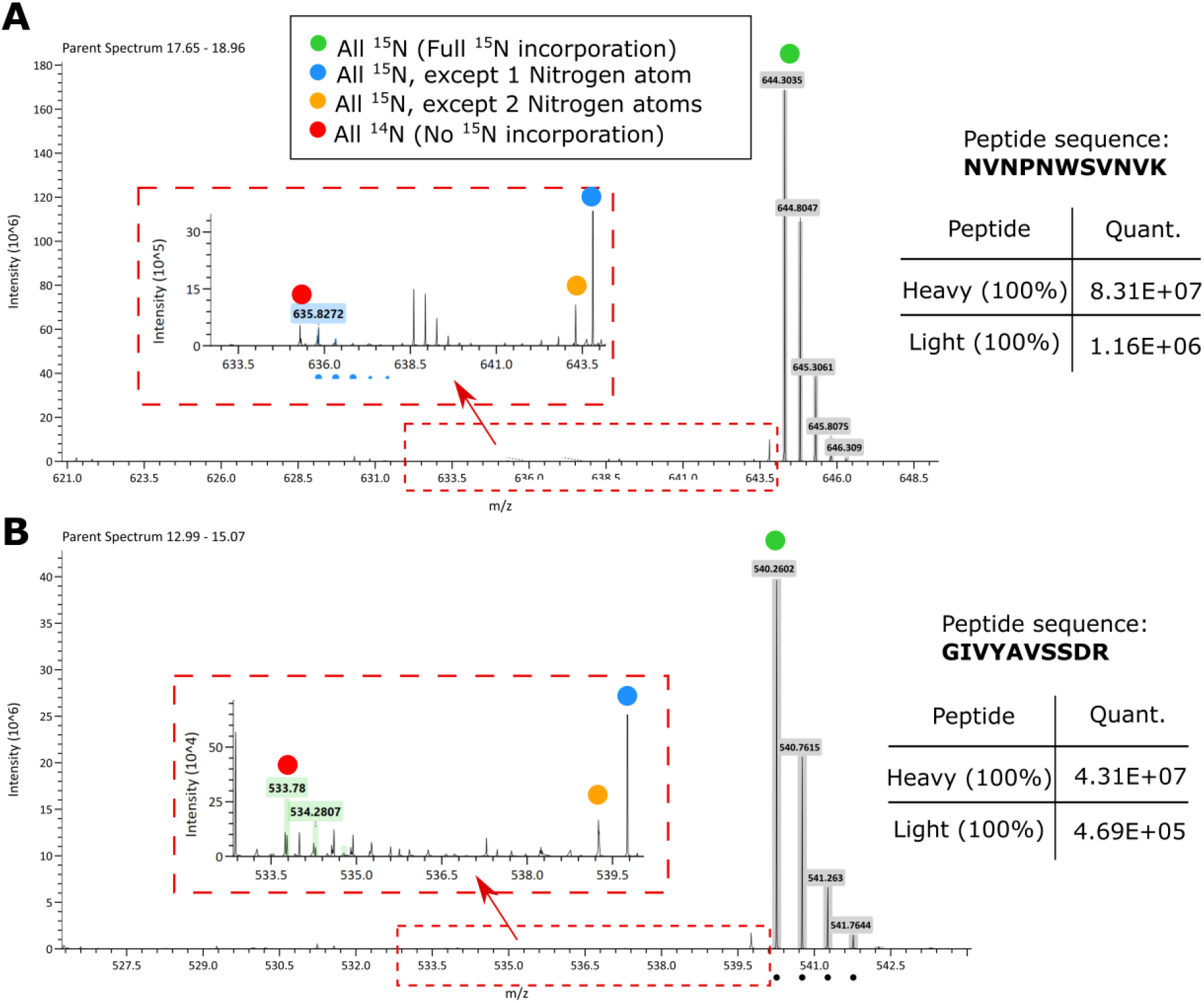
^15^N incorporation in purified heavy labeled DCX was assessed by LC-MS analysis. The two MS1 scans for two candidate DCX peptides are shown as the examples. The highlighted mass to charge ratios (M/Zs) are corresponding to a range of species from fully ^14^N to fully ^15^N species. The corresponding sample area for light and heavy species is shown in the inserted tables.

**Fig. S6.**
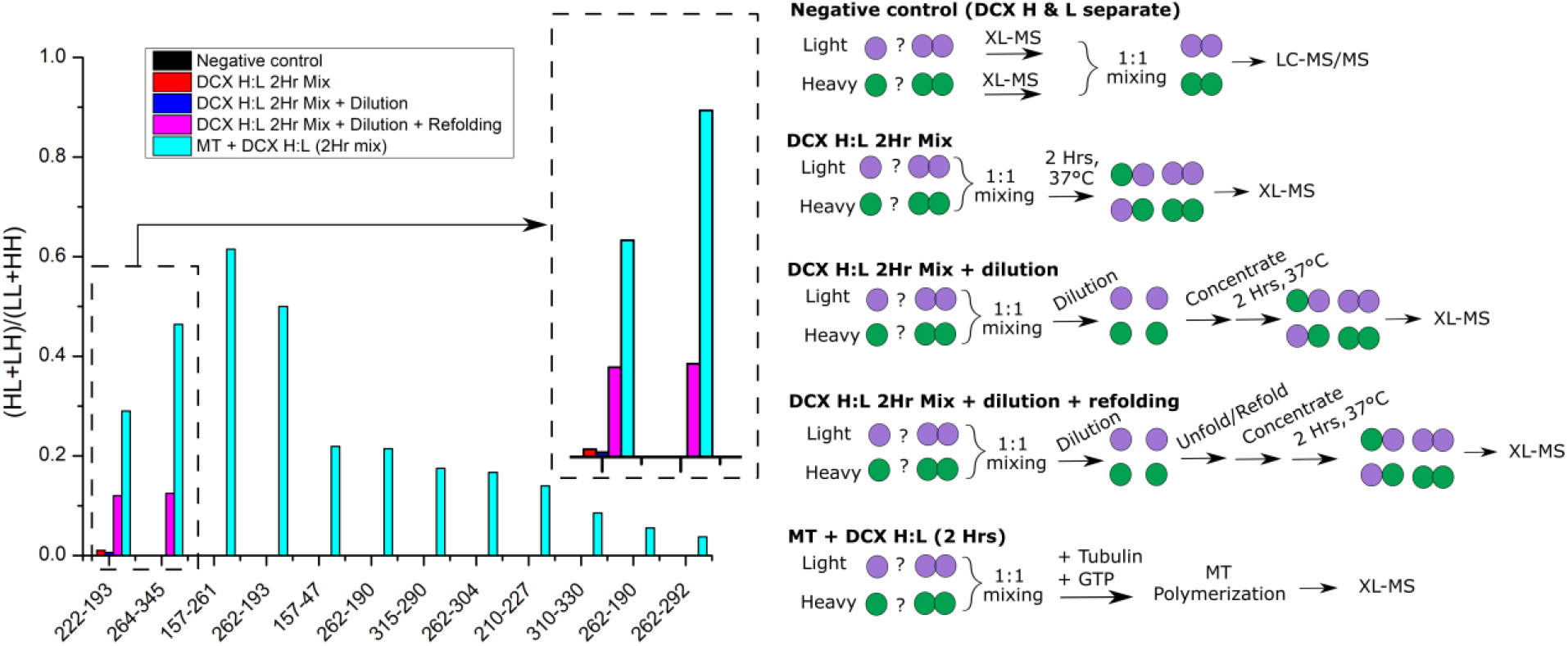
Monitoring the ratio of heterogenous species (Heavy-Light:HL and Light-Heavy:LH) to the homogenous species (Light-Light:LL and Heavy-Heavy:HH), for different crosslinked peptides. The X-axis IDs are crosslinked residues. The experimental procedure for each sample is schematically represented in right. Only two peptides were shared among different samples (221-193 an 264-345). The two peptides were also observed in the H/L sample (negative control) but no HL or LH species were observed.

**Fig. S7-A to -D.**
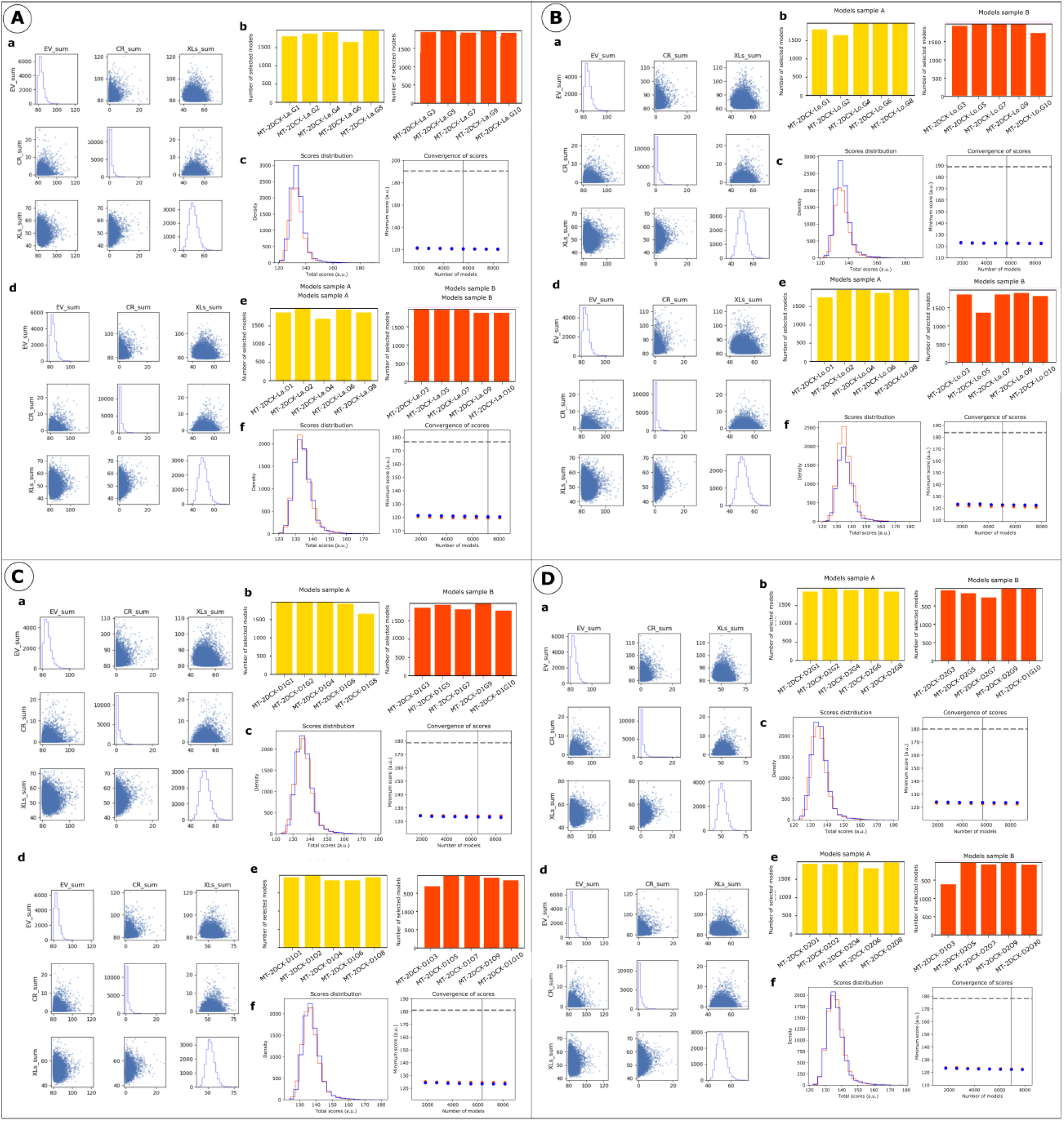
IMP analysis output for MT-dimeric DCX using DCX-DCX crosslinking data, where the NDCs were fixed at lateral (A), longitudinal (B), diagonal 1 (C) and diagonal 2 (D) relative positions on the MT lattice. **a)** The distribution of scores for the models computed for MT-dimeric DCX using the globular CDC structure (100,000 models). **b)** The number of models randomly selected for model samples A (yellow) and B (red), from 10 different modeling runs. **c)** The score convergence for the models in sample A and B. **d)** The distribution of scores for the models computed for MT-dimeric DCX using the open CDC structure (100,000 models). **e)** The number of models randomly selected for model samples A (yellow) and sample B (red), from 10 different modeling runs. **f)** The score convergence for models in sample A and B. The nonparametric Kolmogorov–Smirnov two-sample test (two-sided) indicates that the difference between the two score distributions is insignificant for each modeling run, as the magnitude of the difference is small, demonstrated by the Kolmogorov–Smirnov two-sample test statistic for each modeling run (lateral-globular: 0.010, lateral-open: 0.013, longitudinal-globular:0.014, longitudinal-open: 0.011, diagonal1-globular:0.023, diagonal1-open:0.015, diagonal 2-globular:0.019, and diagonal 2-open:0.014.) Thus, the two score distributions are effectively equal for each modeling run.

**Fig. S8-A to -D.**
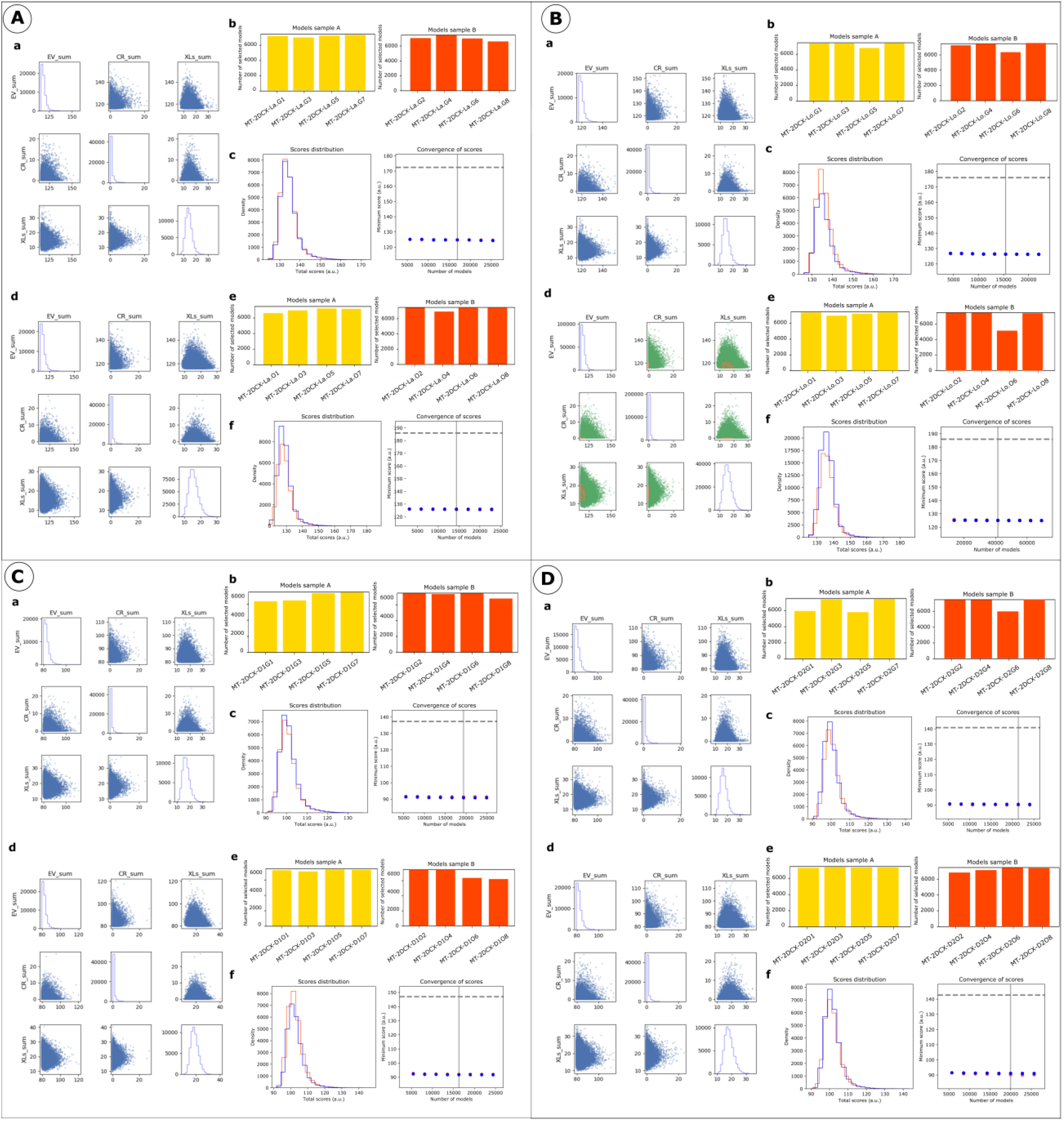
IMP analysis output for MT-dimeric DCX using MT-DCX and DCX-DCX crosslinking data, where the NDCs were fixed at lateral (A), longitudinal (B), diagonal 1 (C) and diagonal 2 (D) relative positions on MT lattice. **a)** The distribution of scores for the models computed for the MT-dimeric DCX using globular CDC structure (320,000 models). **b)** The number of models randomly selected for model samples A (yellow) and B (red), from 10 different modeling runs. **c)** The score convergence for models in sample A and B. **d)** The distribution of scores for all the models computed for MT-dimeric DCX using the open CDC structure (320,000 models). **e)** The number of models randomly selected for model samples A (yellow) and B (red), from 10 different modeling runs. **f)** The score convergence for models in samples A and B. The nonparametric Kolmogorov–Smirnov two-sample test (two-sided) indicates that the difference between the two score distributions is insignificant for each modeling run, as the magnitude of the difference is small, demonstrated by the Kolmogorov–Smirnov two-sample test statistic for each modeling run (lateral-globular: 0.010, lateral-open: 0.010, longitudinal-globular:0.009, longitudinal-open:0.006, diagonal1-globular: 0.007, diagonal1-open:0.010, diagonal 2-globular:0.016, and diagonal 2-open: 0.010). Thus, the two score distributions are effectively equal for each modeling run.

**Fig S9.**
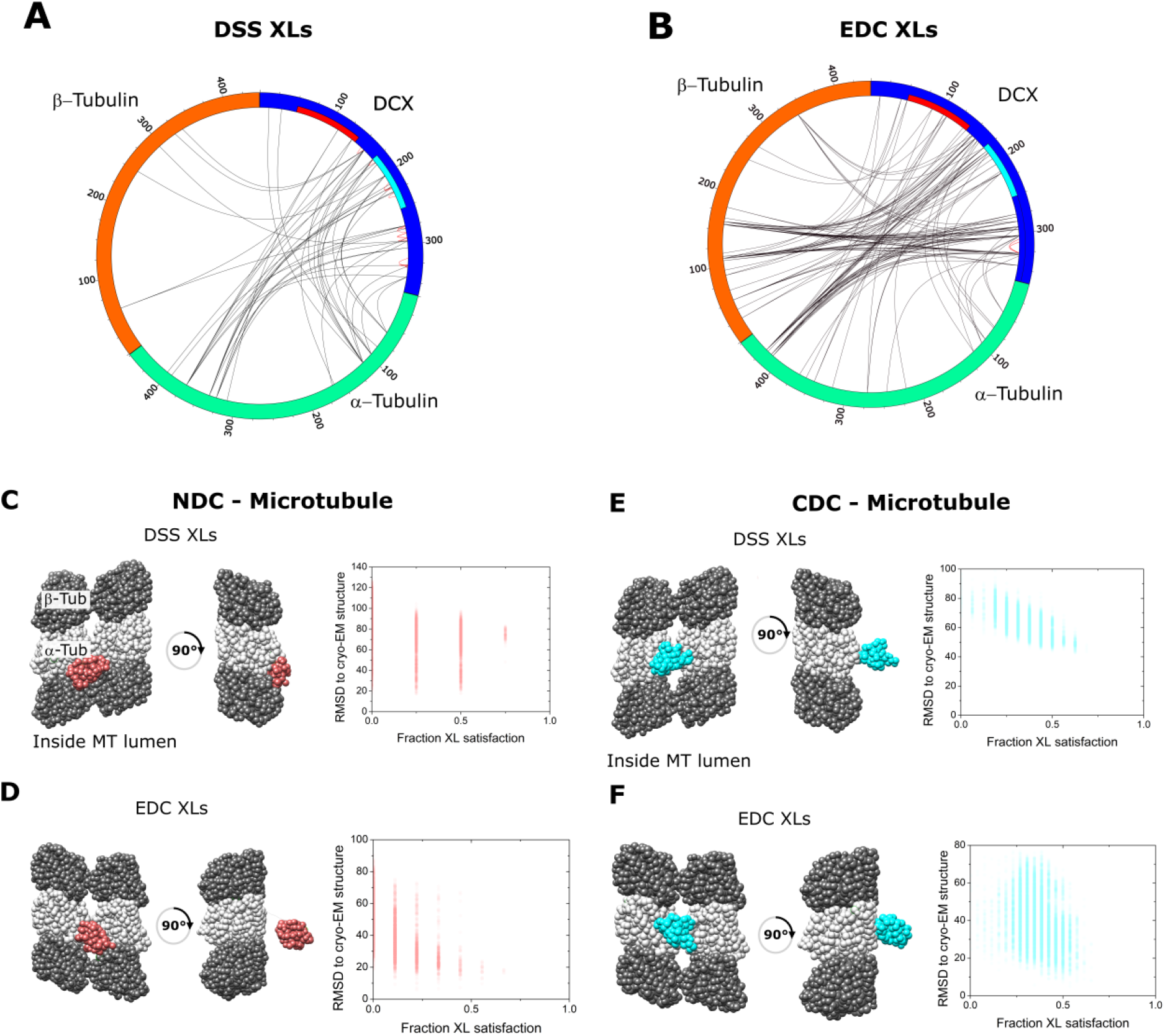
Crosslinking-mass spectrometry analysis of DCX-MT using conventional crosslinkers. Two-dimensional crosslinking map linking α−β Tubulin and DCX at specific residues using (A) DSS and (B) EDC. NDC and CDC domains of DCX sequence are shown with red and cyan, respectively. A subset of inter-DCX crosslinking sites observed among peptides with a shared sequence are shown in red loops. The crosslinking map is produced using xVis online tool [10]. The centroid structure for the major cluster of models produced by IMP for NDC-MT guided exclusively by XL-MS restrains produced by DSS crosslinker (C) and EDC (D). The centroid structure for the major cluster of models produced by IMP for CDC-MT guided exclusively by XL-MS restrains produced by DSS crosslinker (E) and EDC (F); α−Tubulin and β−Tubulin are shown as light and dark grey, respectively. NDC is shown in red, and CDC is shown in cyan. The fraction crosslink satisfaction (defined as <35 Å) versus RMSD to canonical binding site (PDB 4ATU) for all models present in the main structural cluster is presented for each modeling scenarios.

**Fig S10.**
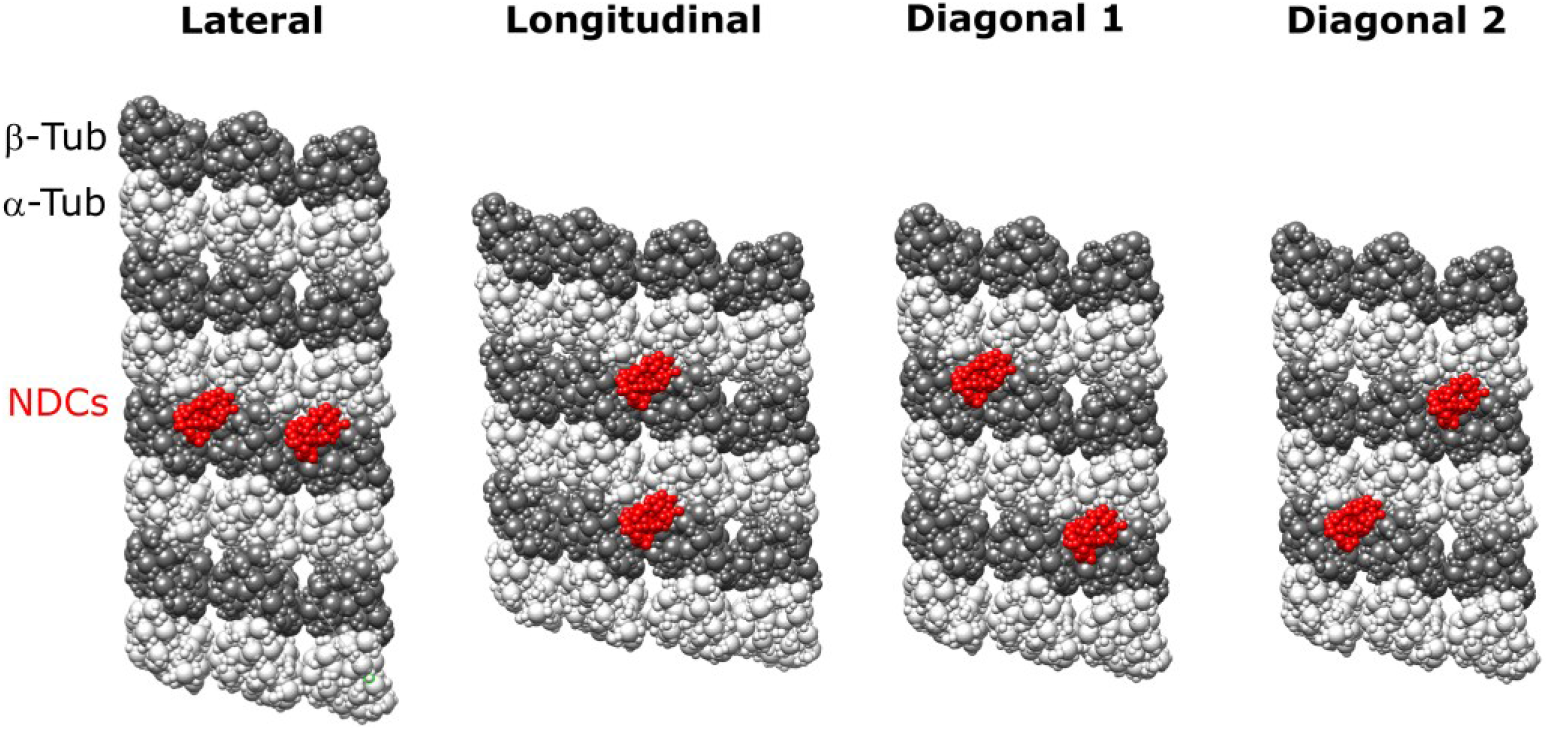
The four different relative positions of fixed NDC on MT lattice that was assessed for ISM of dimeric DCX-MT: Lateral, Longitudinal, Diagonal-1 and Diagonal-2. α−Tubulin and β−Tubulin are shown as light and dark grey, respectively. NDC is shown in red,

